# The myokine FGF21 associates with enhanced survival in ALS and mitigates stress-induced cytotoxicity

**DOI:** 10.1101/2024.09.11.611693

**Authors:** Abhishek Guha, Ying Si, Reed Smith, Mohamed Kazamel, Nan Jiang, Katherine A. Smith, Anna Thalacker-Mercer, Brijesh K. Singh, Ritchie Ho, Shaida A Andrabi, Joao D Tavares Da Silva Pereira, Juliana S. Salgado, Manasi Agrawal, Emina Horvat Velic, Peter H. King

## Abstract

Amyotrophic lateral sclerosis (ALS) is an age-related and fatal neurodegenerative disease characterized by progressive muscle weakness. There is marked heterogeneity in clinical presentation, progression, and pathophysiology with only modest treatments to slow disease progression. Molecular markers that provide insight into this heterogeneity are crucial for clinical management and identification of new therapeutic targets. In a prior muscle miRNA sequencing investigation, we identified altered FGF pathways in ALS muscle, leading us to investigate FGF21. We analyzed human ALS muscle biopsy samples and found a large increase in FGF21 expression with localization to atrophic myofibers and surrounding endomysium. A concomitant increase in FGF21 was detected in ALS spinal cords which correlated with muscle levels. FGF21 was increased in the SOD1^G93A^ mouse beginning in presymptomatic stages. In parallel, there was dysregulation of the co-receptor, β-Klotho. Plasma FGF21 levels were increased and high levels correlated with slower disease progression, prolonged survival, and increased body mass index. In NSC-34 motor neurons and C2C12 muscle cells expressing SOD1^G93A^ or exposed to oxidative stress, ectopic FGF21 mitigated loss of cell viability. In summary, FGF21 is a novel biomarker in ALS that correlates with slower disease progression and exerts trophic effects under conditions of cellular stress.

## Introduction

Amyotrophic lateral sclerosis (ALS) is a multisystemic and age-related neurodegenerative disease characterized by progressive weakness from degeneration of the motor axis including upper and lower motor neurons. This weakness ultimately leads to respiratory failure and death approximately 3 to 5 years after onset.^1,2^ The neuromuscular component of the motor axis plays a pivotal role in progression of ALS, with some investigators postulating a direct role in disease initiation.^3,4^ Indeed, insights from animal studies reveal that initial pathological changes in ALS manifest peripherally, characterized by the breakdown of the neuromuscular junction (NMJ) and the presence of mitochondrial abnormalities within skeletal muscle.^5,6^ We and others have characterized a dysregulated transcriptome in skeletal muscle of ALS patients (see recent review^4^). Our group has identified aberrant expression of proteins and non-coding RNAs reflecting different signaling pathways such as TGFβ/Smad, Wnt, vitamin D and FGF23.^7–9,10,11–13^ Interestingly, many of these molecular patterns overlap in the SOD1^G93A^ mouse, often starting in the early pre-clinical stages and changing with disease progression.^4^ Some of the biomarkers identified contribute to an intricate regulatory network, facilitating molecular crosstalk between motor neurons and myofibers to support the maintenance and regenerative capacity of the neuromuscular unit. On the other hand, other factors may be detrimental to the motor axis.^3,4,14^ Hence, characterizing these biomarkers and exploring the molecular pathways they reflect can lead to improved patient management, including clinical trials, while also identifying novel targets for therapeutic development.^15^

In a prior miRNA sequencing analysis with ALS muscle tissue, we found patterns predicted to alter FGF21 signaling.^12^ Based on this background, we hypothesized that FGF21 may be dysregulated in ALS skeletal muscle early on and serve as a biomarker. FGF21 is a hormone primarily produced in the liver under physiological states and plays a major role in regulating glucose and energy metabolism.^16^ In stressed or pathophysiological conditions, such as mitochondrial stress/dysfunction or endoplasmic (ER) stress, FGF21 can be induced in many tissues, including skeletal and cardiac muscle, where it can mitigate metabolic dysfunction and restore and/or maintain mitochondrial homeostasis through autocrine and paracrine pathways.^17–20^

Here, we utilized an extensive human repository of ALS tissue to investigate FGF21 and found a significant increase of FGF21 expression in ALS muscle, predominantly in atrophied myofibers and surrounding endomysial connective tissue. We found a concomitant increase in the mRNA of the co-receptor, β-Klotho (*KLB*), suggesting that FGF21 signaling is increased in the neuromuscular unit. FGF21 was increased in plasma samples of ALS patients, and high levels associated with slower disease progression and longer survival. Using *in cellulo* models of motor neurons, we found evidence suggesting that FGF21/KLB is dysregulated in ALS motor neurons. FGF21 is altered in both motor neurons and muscle cells with oxidative stress or co-expression of SOD1^G93A^, and it exerts a cytoprotective effect when ectopically expressed.

## Materials and methods

### Human Clinical Samples Collection

Muscle samples and post-mortem tissues were obtained under UAB Institutional Review Board (IRB)-approved protocols as described previously.^7,8,21^ Some of the post-mortem spinal cord samples (normal and ALS) were procured from the Department of Veterans Affairs Biorepository Brain Bank.

Blood samples were collected from non-fasting ALS patients and normal controls enrolled in a prior biomarker study approved by the UAB IRB.^21^ Samples were drawn in EDTA plasma collection tubes, centrifuged at 1600 RCF for 10 min at 4° C, and stored at −80° C. For correlation studies with disease progression and survival, enrolled patients had a sample at study entry and 3 months later.

### Animal Sample Collection

All animal procedures were approved by the UAB Institutional Animal Care and Use Committee and were carried out in accordance with relevant guidelines and regulations of the National Research Council Guide for the Care and Use of Laboratory Animals and in compliance with the ARRIVE guidelines. B6.Cg-Tg (SOD1_G93A) 1 Gur/J male mice (Jackson Laboratory) were generated as previously detailed.^8^ After euthanasia, gastrocnemius muscle and spinal cord tissue samples were collected from SOD1^G93A^ and WT-SOD1 littermate controls at post-natal day 20, 40, 60, 125 and 150. These time points cover the full range of disease stages in the ALS mouse as previously described.^11^

### Cell Culture, Treatment, and Transfection

C2C12 myoblasts (ATCC) and NSC-34 cells were grown in 0.22 μm filter (Corning) sterilized Dulbecco’s modified Eagle’s High Glucose (Corning) pH 7.4, supplemented with 10% FBS (Thermo Fisher Scientific), 1% PEN-STREP (Thermo Fisher Scientific) and 1% L-Glutamine at 37°C in a humidified 5% CO2 incubator. Cells were treated with methionine-cystine (MetCys)-deprived media (Thermo Fisher Scientific) or 100 ng/ml mouse recombinant FGF21 (GenScript) or exposed to 100 µM H_2_0_2_ (Sigma Aldrich) for 24 h. Additionally, some cells were transfected with pCDNA3.1 FGF21-Flag (GenScript), Flag-SOD1^G93A^ and Flag-SOD1WT plasmid constructs^22^ using Lipofectamine 2000 (Thermo Fisher Scientific) in Opti-MEM Reduced Serum Media (Thermo Fisher Scientific). To induce C2C12 myoblast differentiation, growth media (GM) was switched to differentiation media (DM) containing 2% horse serum.

### iPSC-derived Motor Neurons

iPSCs were derived from ALS subjects and controls and are summarized in Supplementary Table 1. Differentiation of motor neurons followed methods previously published by our group^11^ except for the ALS lines, NN0004306 and NN0004307, and the control lines, FA0000011 and NN0003920. For these four lines, a different protocol was used and detailed elsewhere.^23,24^

### Cell proliferation and cell Viability assay

C2C12 cells were transfected with a FLAG-tagged-FGF21 expressing plasmid in a dose-dependent manner, and after 48 h post-transfection, cells were (1 x 10^4^) seeded in 96-well plate. Cell proliferation was assessed at intervals of 24, 48 and 72 h using the MTS-based cell growth determination kit (Promega). For oxidative stress, cells were incubated with MetCys-deprived media or H_2_O_2_ treatment. Cell viability was determined with the ViaLight Plus BioAssay kit (Lonza) following the manufacturer’s instructions and as previously described.^25^ Apoptosis was detected by caspase activity using Caspase-Glo 3/7 assay kit (Promega) following the manufacturer’s protocol.

### RNA analysis

RNA was extracted from muscle and spinal cord tissue samples using TRIzol (Thermo Fisher) and RNASpin Mini kit (Cytiva) per the manufacturer’s instructions. RNA from iPSC motor neurons was extracted using the RNeasy kit (Qiagen) following the manufacturer’s instructions. For other cell culture samples, cells were lysed using the lysis buffer provided in the RNASpin Mini kit. cDNAs were synthesized using the High-Capacity cDNA Reverse Transcription Kit (Thermo Fisher Scientific). mRNA expression of *FGF21*, and *KLB*, was quantified utilizing Taqman primers (Thermo Fisher) with the QuantStudio 5 Real-Time PCR system (Thermo Fisher). *RPS9* and *TBP1* were used as internal controls. For iPSC-derived motor neurons, a fold-change in RNA for ALS motor neurons was generated by comparing to control lines within the same differentiation protocol.

### Immunoblotting and Immunochemistry

Immunoblotting was performed based on methods described previously.^8^ The following antibodies were used: anti-FGF21 (Abcam), 1:1000; anti-MHC (Developmental Studies Hybridoma Bank), 1:200; anti-FLAG-M2 (Sigma Aldrich), 1:1000; anti-Vinculin (Abcam), 1:3000.

For immunocytochemistry, 1 x 10^5^ C2C12 cells were seeded in Nunc Chamber Slide System (Thermo Fisher) and then treated with DM 24 h later. Cells were fixed at different time intervals and blocked using Intercept (PBS) Blocking Buffer (Li-COR) for 30 min. Subsequently, cells were stained using an anti-MHC antibody (Developmental Studies Hybridoma Bank), 1:50, overnight at 4° C, followed by an anti-Mouse Alexa Fluor 488 secondary antibody (Thermo Fisher), 1:400. Nuclei were stained with DAPI (Thermo Fisher) at 1:2000 for 5 min at RT. Images were captured using a Nikon C2 confocal microscope. The fusion index was calculated as the number of nuclei inside MHC-positive myotubes divided by the total number of nuclei present in a field of view from three random fields per biological replicate (total of 3 replicates).^26^

For immunohistochemistry, muscle samples were fixed in 4% PFA at 4°C overnight and embedded in paraffin. Deparaffinized 10 µm sections were boiled in 10mM citrate buffer (pH 6.0) for 30 min, and cooled to RT. Slides were incubated in 3% H_2_O_2_ for 10 min and then incubated with FGF21 antibody (Abcam), 1:100, overnight at 4°C. After washing in PBS, slides were incubated with a donkey anti-rabbit Cy3 secondary antibody (Jackson Immunoresearch) for 90 min at RT. Sections were incubated in Wheat Germ Agglutinin (WGA), Oregon Green 488 Conjugate (Thermo Fisher), followed by Hoechst 33342 (Sigma Aldrich) at 1:20,000 for 5 min. Confocal imaging was done to assess atrophic and non-atrophic muscle fibers. Mean FGF21 fluorescence intensity (MFI) was assessed on muscle tissue samples as described previously.^7,21^ The minimal Feret’s diameter of selected myofibers determined myofiber size, with fibers less than 25 μm in diameter considered atrophic. A total of 46 atrophic fibers were compared to an equal number of non-atrophic fibers within the same images.

### ELISA

FGF-21 was quantitated in muscle, spinal cord and plasma samples using U-PLEX Human FGF-21 ELISA Assay (K1515WK-1, MSD) according to the manufacturer’s instruction. For survival and rate of disease progression, the FGF21 level at study entry and the 3-month time interval were averaged. To evaluate the levels of secreted FGF21 protein, conditioned media (CM) were collected from C2C12 and NSC-34 cells following exposure to different treatments or conditions. FGF21 levels were quantitated using U-PLUX mouse-FGF21 Assay (K1525WK-1, MSD).

### Statistics

Statistical analyses for both human and mouse data were conducted using GraphPad Prism 10.2 (San Diego, CA, USA). *FGF21* mRNA expression in human muscle biopsy samples was analyzed with a two-tailed t-test with a Welch’s correction. For *KLB* mRNA, a two-tailed Mann Whitney test was used for comparison. A two-tailed Mann Whitney test was used for post-mortem tissues, FGF21 quantitative immunostaining, and iPSC-derived motor neurons. A two-tailed t-test was used for mouse muscle and spinal cord tissue and cell culture analyses, plasma FGF21, BMI, viability, and Caspase-3/7 activity. One-way ANOVA with Tukey’s multiple comparisons test was used for comparisons of plasma FGF21 levels between different disease progression subsets, cell stress-induced changes in viability, and C2C12 proliferation assays. A Spearman correlation test was used for spinal cord versus muscle FGF21 protein levels, muscle *FGF21* versus *HDAC4* mRNA levels, and ΔALSFRS versus plasma FGF21 levels. A Log-rank (Mantel-Cox) test was used to compare Kaplan Meier survival curves in ALS patients.

## Results

### FGF21 is increased in human ALS muscle and spinal cord tissue

We assessed FGF21 mRNA expression in a large cohort of muscle biopsy samples of ALS patients from our clinic (Supplementary Table 2 and Fig. 1). Demographics for this cohort, including age (mean 57 ± 13 y) and slight male to female predominance (1.4:1), were in line with prior epidemiological studies.^1,27^ The cohort of normal biopsies showed similar age with a slight female to male predominance (1.2:1). By qPCR, we found an ∼8-fold increase in *FGF21* mRNA in ALS samples (*P =* 0.004) compared to normal control biopsy samples (Fig. 1A). There was variability in mRNA levels with several at greater than 50-fold higher than controls. We next tested post-mortem ALS muscle samples which reflect more advanced (end-stage) disease as reflected by the longer disease duration (51 months versus 15 months in the biopsy group). *FGF21* mRNA levels were much higher at 63-fold greater than normal control samples, indicating that *FGF21* expression in muscle increases with disease progression (Fig. 1B). We compared *FGF21* mRNA levels with *HDAC4*, another known marker of muscle denervation in ALS,^28^ and found suggestive evidence of a correlation (Spearman rank correlation *R* = 0.683, *P* = 0.050, Supplementary Fig. 1). We next assessed FGF21 protein expression by ELISA with post-mortem samples and found a ∼7-fold increase in ALS samples (2463 versus 335 pg/mg of muscle tissue; *P =* 0.030). As with mRNA levels, there was variability among ALS samples (versus controls) with one ALS patient showing nearly 15,000 pg/mg. We next measured *FGF21* mRNA levels in spinal cord tissue but observed no significant difference compared to control tissue as a whole; however, there were 4 outliers with markedly increased levels of up to 25-fold (Fig. 1C). On the other hand, FGF21 protein was increased in ALS spinal cord (132 versus 29 pg/mg of spinal cord tissue; *P =* 0.005), but levels in spinal cord were nearly 19-fold less than ALS muscle. There was variability among the ALS patients as observed with muscle. Post-mortem spinal cord and muscle samples were matched for 18 of the ALS patients tested and they showed a positive correlation for FGF21 levels (Spearman rank correlation *R* = 0.49, *P =* 0.039) (Fig. 1D). We next sought to determine whether changes in *FGF21* occurred in the SOD1^G93A^ mouse model of ALS. This model can provide insight into temporal patterns of biomarkers identified in human ALS muscle including pre-clinical (prodromal) phases of the disease.^4^ We detected a 4-fold increase in *FGF21* mRNA compared to age-matched wild-type (WT) controls at 40 d post-natal (Fig. 1E). Although NMJ innervation is reduced by 40% by this age, this stage is typically considered pre-symptomatic where standard testing such as rotarod and grip strength are unaffected and only subtle signs of muscle weakness can be detected by more in-depth testing.^29^ In later stages, muscle FGF21 rose from ∼37-fold at 60 d (pre-symptomatic) to 43-fold at 125 d (symptomatic stage). At end-stage (150 d), there was a 28-fold fold-change. FGF21 mRNA increased in spinal cord tissue in parallel with muscle FGF21. Taken together, FGF21 mRNA and protein levels were significantly but variably increased in human ALS muscle and spinal cord tissues, with a disproportionate increase in muscle. Furthermore, the strong association between muscle and spinal cord FGF21 for each patient and in the SOD1^G93A^ mouse model suggested an underlying connectivity.

**Figure 1.**
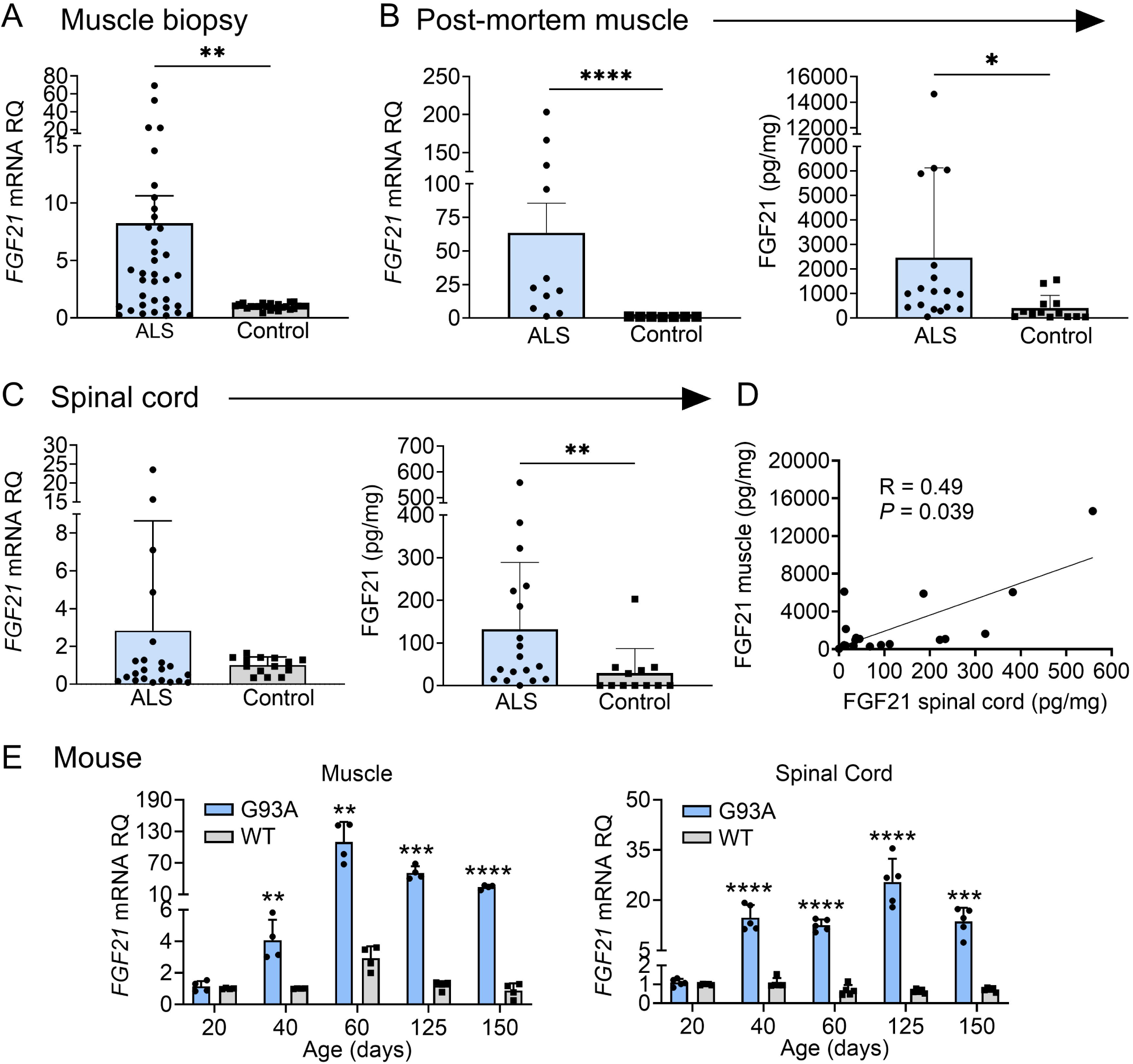
FGF21 levels are elevated in the muscle and spinal cord tissues of ALS patients and the SOD1^G93A^ mouse. (**A**) *FGF21* mRNA expression was analysed in normal (*n* = 24) and ALS (*n* = 36) muscle biopsy samples via RT-qPCR. \*\**P =* 0.004, unpaired two-tailed t-test with Welch’s correction. (**B**) *FGF21* mRNA levels were quantified in normal (*n* = 7) and ALS (*n* = 11) post-mortem muscle samples (left panel) and FGF21 protein levels (*n* = 13 for normal samples; *n* = 18 for ALS samples; right panel). \*\**P =* 0.003, \*\*\*\**P <* 0.0001, two-tailed Mann Whitney test. **(C)** *FGF21* mRNA levels were quantified in normal (*n* = 14) and ALS (*n* = 22) post-mortem spinal cord samples (left panel) and FGF21 protein levels (*n* = 12 for normal samples; *n* = 18 for ALS samples; right panel). \*\**P =* 0.00, two-tailed Mann Whitney test. **(D)** Comparison of spinal cord and muscle FGF21 protein levels for 18 ALS patients. A spearman correlation test was used for analysis. **(E)** *FGF21* mRNA levels in the gastrocnemius muscle (left panel) and spinal cord (right panel) were quantified across different age groups (20 – 150 days; *n* = 4-5 per group) from SOD1^G93A^ mice and littermate controls. \*\**P <* 0.01, \*\*\**P <* 0.001, \*\*\*\**P <* 0.0001, unpaired two-tailed t-test comparing WT to SOD1^G93A^. For all graphs, error bars represent SD.

### FGF21 localizes to atrophic muscle fibers

Next, we determined localization of FGF21 in ALS muscle by immunohistochemistry. We observed intense intra-myofiber staining in areas of grouped atrophy (characterized by clusters of small angular fibers) which is a hallmark of denervation (Fig. 2A).^30^ FGF21 staining also localized to the endomysial connective tissue. On the other hand, non-atrophic fibers within the same section showed little or no staining. Five normal control muscle samples also showed no FGF21 staining (example shown in Fig. 2A). To confirm an association between FGF21 immunoreactivity and atrophic myofibers, we measured mean fluorescence intensity (MFI) in atrophic and non-atrophic fibers in muscle sections from five ALS patients (Fig. 2B). Regions of interest (ROI) for atrophic and non-atrophic myofibers were selected within the same muscle section, often in proximity, to minimize the effects of variable tissue staining (see example in Fig. 2B). We observed a 9-fold increase in FGF21 MFI in ROIs associated with atrophic versus non-atrophic myofibers (*P* < 0.0001). Taken together, these data indicate a marked increase in FGF21 expression in ALS muscle tissues, predominantly in atrophic myofibers and surrounding endomysial connective tissue.

**Figure 2.**
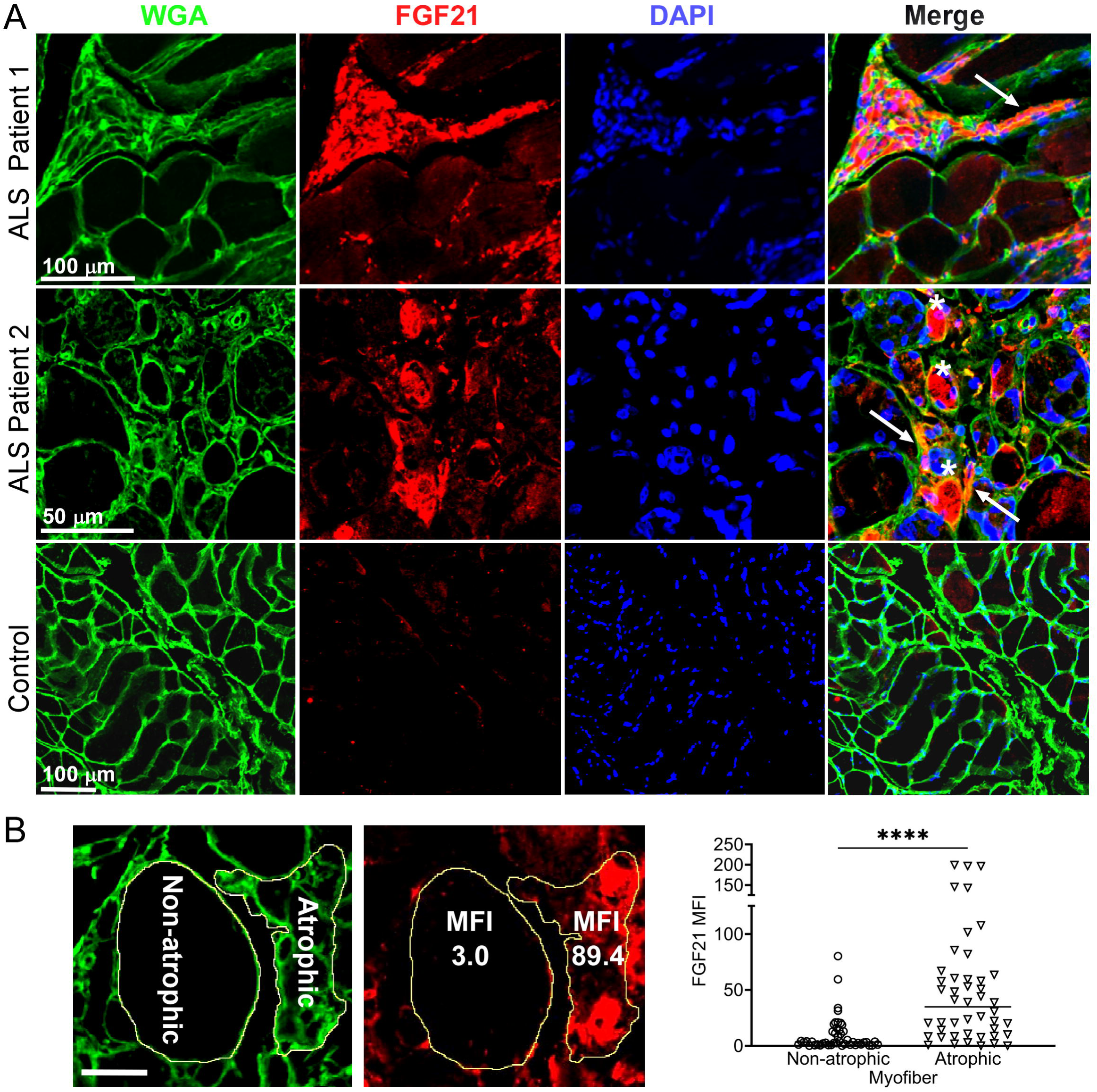
FGF21 localizes to atrophic myofibers in ALS muscle. **(A)** Tissue sections from two ALS patients and one normal control were immunostained with an anti-FGF21 antibody and counterstained with Hoechst and wheat germ agglutin (WGA). Intense immunoreactivity is observed in atrophic myofibers (asterisks) and in the endomysial space (arrows) in the ALS muscle sections. Scale bars, 100 µM in low power views and 50 µM in the enlarged views. **(B)** Mean fluorescence Intensity (MFI) analysis of FGF21 immunoreactivity was performed in 5 ALS patient biopsy samples and 5 normal controls. Atrophic (< 25 μM minimal Feret’s diameter) and non-atrophic myofibers were selected in the same section as shown in the micrograph (yellow outline). FGF21 MFI (per μM^2^) was quantitated for 46 atrophic and non-atrophic myofibers and summarized in the graph (horizontal line represents the mean). \*\*\*\**P <* 0.0001; two-tailed Mann Whitney test. Scale bar: 50 µM.

### Plasma FGF21 levels are elevated in ALS patients and correlate with enhanced survival

Since FGF21 is a secreted factor and we observed extra-myofiber immunostaining in ALS muscle tissue, we queried whether FGF21 could be detected in plasma of ALS patients. To assess this possibility, we assayed plasma samples collected from a cohort of ALS patients in our clinics who participated in a prior biomarker study (Supplementary Table 3).^9^ A set of age-matched normal controls was used for comparison. Overall, we found a significant increase in FGF21 levels in the ALS cohort (923 v. 649 pg/ml; *P* = 0.03), but with variability (Fig. 3A). Interestingly, the FGF21 levels were more than 2-fold higher in ALS muscle versus plasma FGF21 (albeit end-stage ALS muscle; Fig. 1B). None of the patients or controls had concomitant type II diabetes or known liver disease, two conditions associated with higher circulating levels of FGF21.^31^ We next examined a subset of 16 ALS patients who were followed prospectively with serial examinations and plasma sample collection. Based on calculated monthly changes in ALSFRS-R scores (ΔFRS), there were 6 slow, 5 average and 5 fast progressors using previously established criteria (Fig. 3B).^32^ Using the average of baseline and 3-month FGF21 plasma levels, we found that the slow progressors had more than a 4-fold increase in FGF21 compared to fast progressors (2247 v. 529 pg/ml; P = 0.003) and a ∼3.5-fold increase compared to the control group (P = 0.0003; Fig. 3C). The group of average progressors was intermediate between the slow and fast groups (1410 pg/ml) but did not reach statistical significance. Independent of clinical classification, there was a strong negative correlation between the ΔFRS for each subject and their respective plasma FGF21 level (*R* = −0.710, *P* = 0.003; Fig. 3D). We then assessed survival of these patients based on FGF21 levels, comparing those with <1.5 fold-change (FC) over controls versus ≥ 1.5 FC (Fig. 3E). The median survival for the group with low FGF21 levels was 18 months versus 75 months for the group with high FGF21 levels (*P* = 0.01). Three patients in the high FGF21 group are still alive at 82, 93, and 94 months after disease onset. We next looked at body mass index (BMI) and found that the <1.5 FC group had a significantly lower BMI than the >1.5 FC group (22.6 ± 3.6 v. 29.0 ± 6.6 kg/M^2^, *P* = 0.04; Fig. 3F). Age and duration of disease were not significantly different between the two groups (Supplementary Table 4). Interestingly, in the lower FGF21 group, 4 out of 7 patients had bulbar onset of ALS versus the high FGF21 group which all had spinal onset. Since liver is a major source of FGF21,^33^ we assessed expression in the SOD1^G93A^ mouse at a time point of peak FGF21 expression in skeletal muscle (Supplementary Fig. 2). Similar to results in Fig. 1, we observed a ∼150-fold increase in FGF21 mRNA in the SOD1^G93A^ mouse compared to WT control muscle (set a 1). In liver, the increase was significantly less at 75-fold in the SOD1^G93A^ mouse (*P* < 0.0001) although the mRNA level in WT liver was ∼35-fold greater than WT muscle. Liver tissue was not available in our post-mortem repository to confirm whether this discrepancy applies more broadly to ALS patients. Taken together, had an overall increase in plasma FGF21 and high levels correlated with slower disease progression and longer survival. While FGF21 mRNA is increased in muscle and liver tissues in the SOD1^G93A^ mouse, the disproportionately higher increase in muscle suggests that this tissue is a major contributor to circulating FGF21.

**Figure 3.**
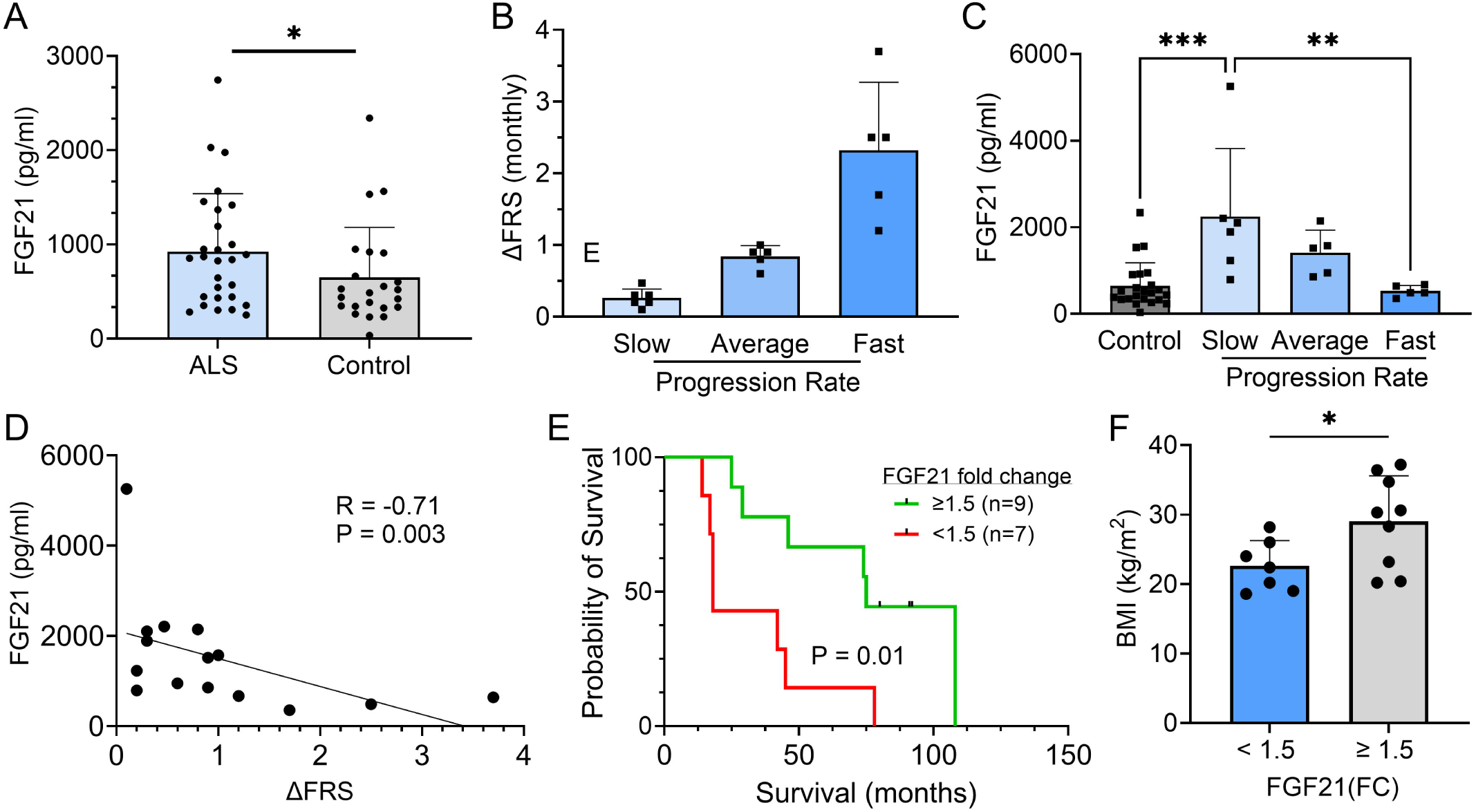
Plasma FGF21 is increased in ALS patients and high levels associate with slower disease progression and prolonged survival. **(A)** Plasma samples from age-matched normal controls (*n* = 23) and ALS patients (*n* = 28) and assayed by ELISA for FGF21. \**P =* 0.043, unpaired two-tailed t-test. **(B)** 16 ALS patients from a prior biomarker study were divided into slow (*n* = 6), average (*n* = 5), and fast (*n* = 5) progressing groups based on the average in the study monthly decline in ALSFRS-R scores. **(C)** Plasma FGF21 levels were measured at baseline and 3 months and averaged. The normal control values were added for comparison. \*\**P =* 0.003, \*\*\**P =* 0.0003; one-way ANOVA followed by Tukey post hoc test. (D) Correlation between plasma FGF21 levels with monthly change in the ALSFRS-R scores (ΔFRS) for each subject. Spearman correlation test. **(E)** Kaplan–Meier survival curves for study patients whose FGF21 plasma levels were < 1.5-fold-change (FC) over the normal control group versus study patients with ≥ 1.5-FC in FGF21 levels. \**P =* 0.015; Log-rank (Mantel-Cox) test. **(F)** Comparison of body mass index (BMI) between the < 1.5 FC and ≥ 1.5-FC groups. \**P =* 0.037; unpaired two-tailed t-test. For all graphs, error bars represent SD.

### FGF21 co-receptor, β-Klotho, is altered in ALS muscle and spinal cord tissue

Given the critical role of β-Klotho (KLB) co-receptor in FGF21 signaling,^33^ we assessed *KLB* mRNA expression in muscle biopsy samples and found a ∼ 4-fold increase in *KLB* levels in the ALS group versus controls (Fig. 4A; *P =* 0.005). On the other hand, post-mortem ALS muscle samples had a 50% reduction in *KLB* mRNA levels (*P =* 0.03) compared to normal controls (Fig. 4B). Likewise, ALS spinal cord tissue showed a significant reduction in *KLB* levels (*P =* 0.0006) compared to controls (Fig. 4C). Some variability was observed with three samples having increased mRNA. Spinal cord tissue from end-stage SOD1^G93A^ mice paralleled these findings and showed a two-fold reduction of *KLB* mRNA compared to littermate controls (*P =* 0.008; Fig. 4D). Collectively, these findings suggest a dysregulation of *KLB* expression in ALS muscle and spinal cord tissues that changed with disease stage and may relate to a progressive loss of motor neurons.

**Figure 4.**
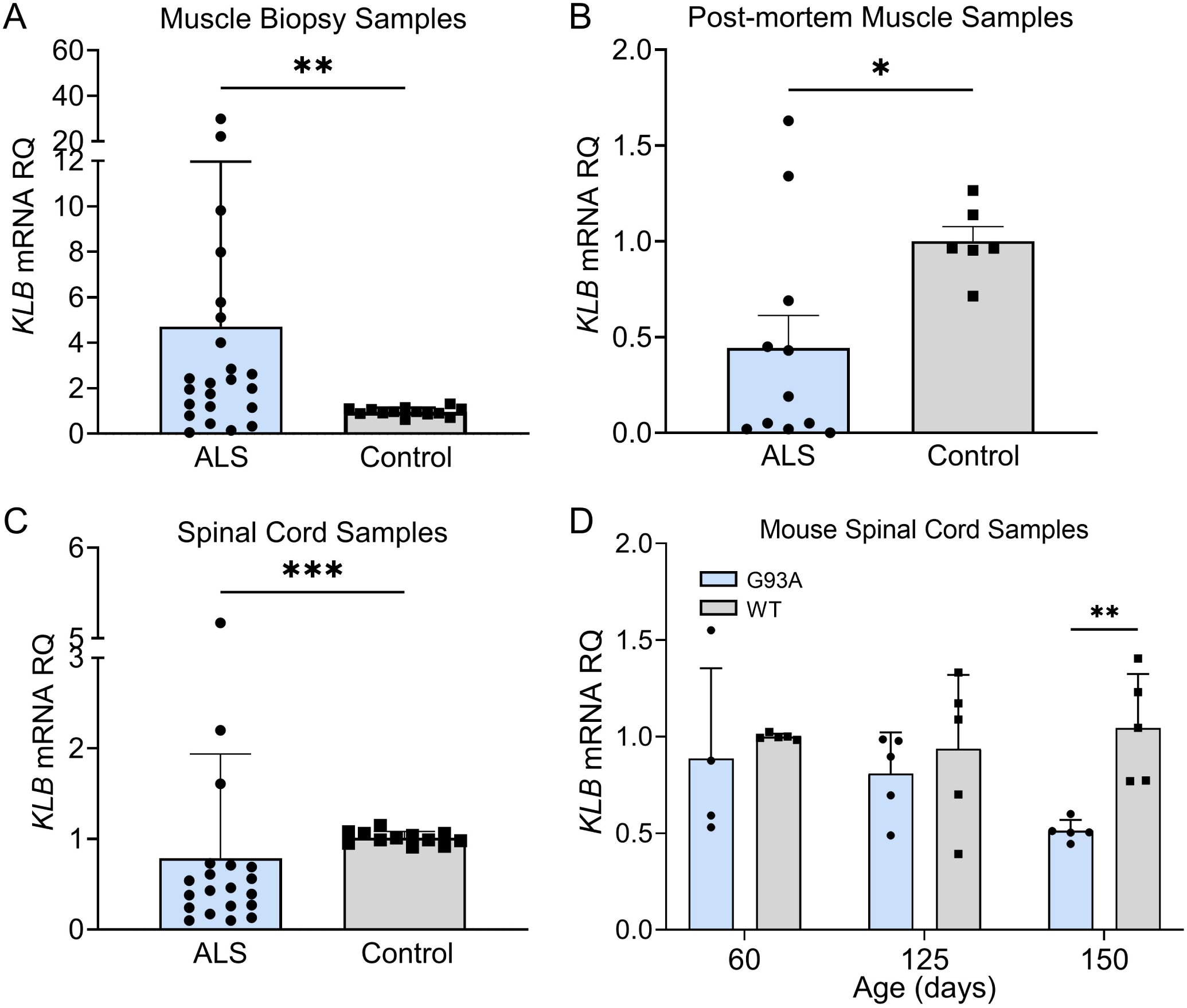
FGF21 coreceptor, *β-Klotho* (*KLB*), is dysregulated in ALS muscle and spinal cord. **(A)** *KLB* mRNA levels were measured in muscle biopsy samples from normal (*n* = 13) and ALS patients (*n* = 23). \*\**P =* 0.005, two-tailed Mann Whitney test. **(B)** *KLB* mRNA levels were measured in post-mortem muscle samples from normal controls (*n* = 6) and ALS (*n* = 11) patients. \**P =* 0.034, two-tailed Mann Whitney test. **(C)** *KLB* mRNA levels were measured in spinal cord samples from normal controls (*n* = 12) and ALS patients (*n* = 20). \*\*\**P =* 0.0006, two-tailed Mann Whitney test. **(D)** *KLB* mRNA levels were measured in spinal cord samples from SOD1^G93A^ mice (*n* = 5) and WT controls (*n* = 5) at different ages. \*\**P =* 0.003; unpaired two-tailed t-test comparing WT to SOD1^G93A^.

### FGF21-KLB axis is dysregulated in ALS motor neurons

As we found progressive loss of *KLB* expression in ALS spinal cord and muscle, we assessed the FGF21-KLB axis in motor neurons. Using iPSC-derived motor neurons from ALS patients (Supplementary Table 1), we assessed the expression of *FGF21* and *KLB* mRNAs. Compared to normal control iPSC motor neurons, there was a ∼50% reduction in *FGF21* mRNA levels in ALS motor neuron samples (Fig. 5A; *P =* 0.012). On the other hand, *KLB* mRNA was increased by 3-fold (Fig. 5B; *P =* 0.018). We next transfected NSC-34 motor neuron like cells with Flag-tagged wild-type (WT) SOD1 or SOD1^G93A^ expression cassettes (Fig. 5C) and found a similar pattern of expression with a more than 2-fold increase in *KLB* (Fig. 5D) and a ∼15% reduction in *FGF21* in SOD1^G93A^-expressing cells versus WT (Fig. 5E). FGF21 protein measured in the CM was unchanged (Fig. 5F). As FGF21-KLB signaling is a major cellular stress response,^16^ we assessed the effect of oxidative stress, on this pathway in NSC-34 cells. When cells were exposed to oxidative stress either by methionine/cysteine (MetCys) deprivation which leads to depletion of the intracellular antioxidant, glutathione,^34,35^ or directly with H_2_O_2_,^34^ there was a marked induction of *FGF21* and *KLB* mRNAs (Fig. 5G and 5H). Induction of these genes was particularly pronounced with MetCys deprivation where *KLB* and *FGF21* mRNAs increased by nearly 100-fold and 25-fold respectively. The induction by H_2_O_2_ was less at two to three-fold. This response was mirrored by induction of stress-response genes, *ATF4* and *PGC-1α*, with MetCys deprivation showing the highest induction (Supplementary Fig. 3A and 3B). *ATF4* was not induced by H_2_O_2_ in NSC-34 cells.

**Figure 5.**
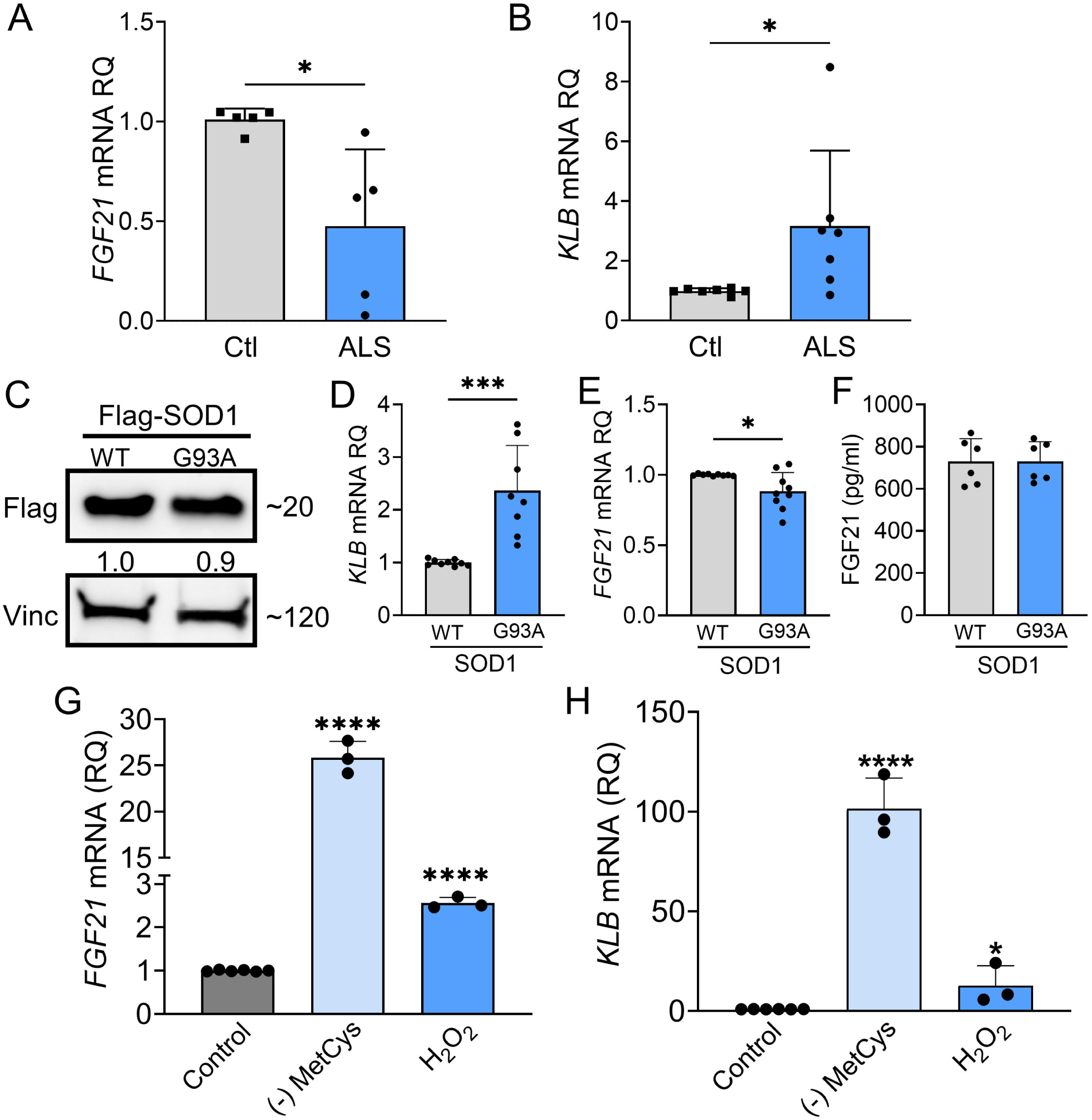
FGF21-KLB signaling is dysregulated in ALS motor neurons. **(A)** *FGF21* were measured in iPSC-derived motor neurons obtained from healthy controls and ALS patients carrying either C9orf72 or SOD1 mutations (Supplementary Table 1). \**P =* 0.012; two-tailed Mann Whitney test. **(B)** *KLB* mRNA levels were similarly measured in iPSC motor neurons. \**P =* 0.018; two-tailed Mann Whitney test. **(C)** NSC-34 motor neuron-like cells were transfected with FLAG-tagged WT and SOD1^G93A^ expression plasmids and lysates were assessed by western blot using the antibodies indicated. Bands were quantitated by densitometry and a ratio to the loading control, vinculin, was calculated (shown between the two blots). **(D)** *KLB* and **(E)** *FGF21* mRNA levels were measured in the same lysates. \**P =* 0.018, \*\*\**P =* 0.0002, unpaired two-tailed t-test. **(F)** FGF21 protein was measured in the CM of transfected NSC-34 cells. **(G)** *FGF21* or **(H)** *KLB* mRNA levels were quantified from NSC-34 cells exposed to methionine-cystine (MetCys)-deprived media or treated with 100μM H_2_O_2_ for 24 h. \**P =* 0.048, \*\*\*\**P <* 0.0001; one-way ANOVA followed by Tukey’s multiple comparisons test. Data points represent biological replicates and bars are the mean ± SD.

### FGF21 mitigates cytotoxic stress in mutant SOD1-expressing NSC-34 cells

We first determined the impact of SOD1^G93A^ on cell viability and found a 33% reduction in viability (Fig. 6A) and a concomitant 50% increase in Caspase-3/7 activity consistent with apoptotic activity (Fig. 6B). Next, we ectopically expressed FGF21 in NSC-34 cells co-expressing wild-type (WT) SOD1 or SOD1^G93A^ to determine the effect on cell viability. We assayed the CM by ELISA and western blot and found a large increase in FGF21 in transfected cells at ∼35 ng/ml (Fig. 6C). Cell viability increased back to control levels along with reduced Caspase-3/7 activity (Fig. 6D and 6E). In a separate experiment, we added recombinant FGF21 to the culture media of SOD1^G93A^-expressing NSC-34 cells and observed a similar reversal of cytotoxicity (Supplementary Fig. 4). To assess further a potential paracrine effect of FGF21, NSC-34 cells expressing either (WT) SOD1 or SOD1^G93A^ and C2C12 cells expressing FGF21-Flag or EV were co-cultured in a transwell plate with a 3 µm porous filter (Supplementary Fig. 5A). Viability was checked and found to be significantly reduced in NSC-34 cells expressing SOD1^G93A^ co-cultured with C2C12-EV cells (*P* < 0.0001, Supplementary Fig. 5B). When co-cultured with C2C12-FGF21 cells, loss of viability was significantly blunted (*P* < 0.0001). We next exposed NSC-34 cells to MetCys deprivation or H_2_O_2_ which led to a significant loss of viability (*P* < 0.0001, Fig. 6F) which was reversed with ectopic FGF21 (Fig. 6G). In summary, *FGF21* and *KLB* mRNAs are expressed in motor neurons and induced by co-expression of ALS-associated SOD1^G93A^ or oxidative stressors, and FGF21 can reverse the resultant loss of cell viability.

**Figure 6.**
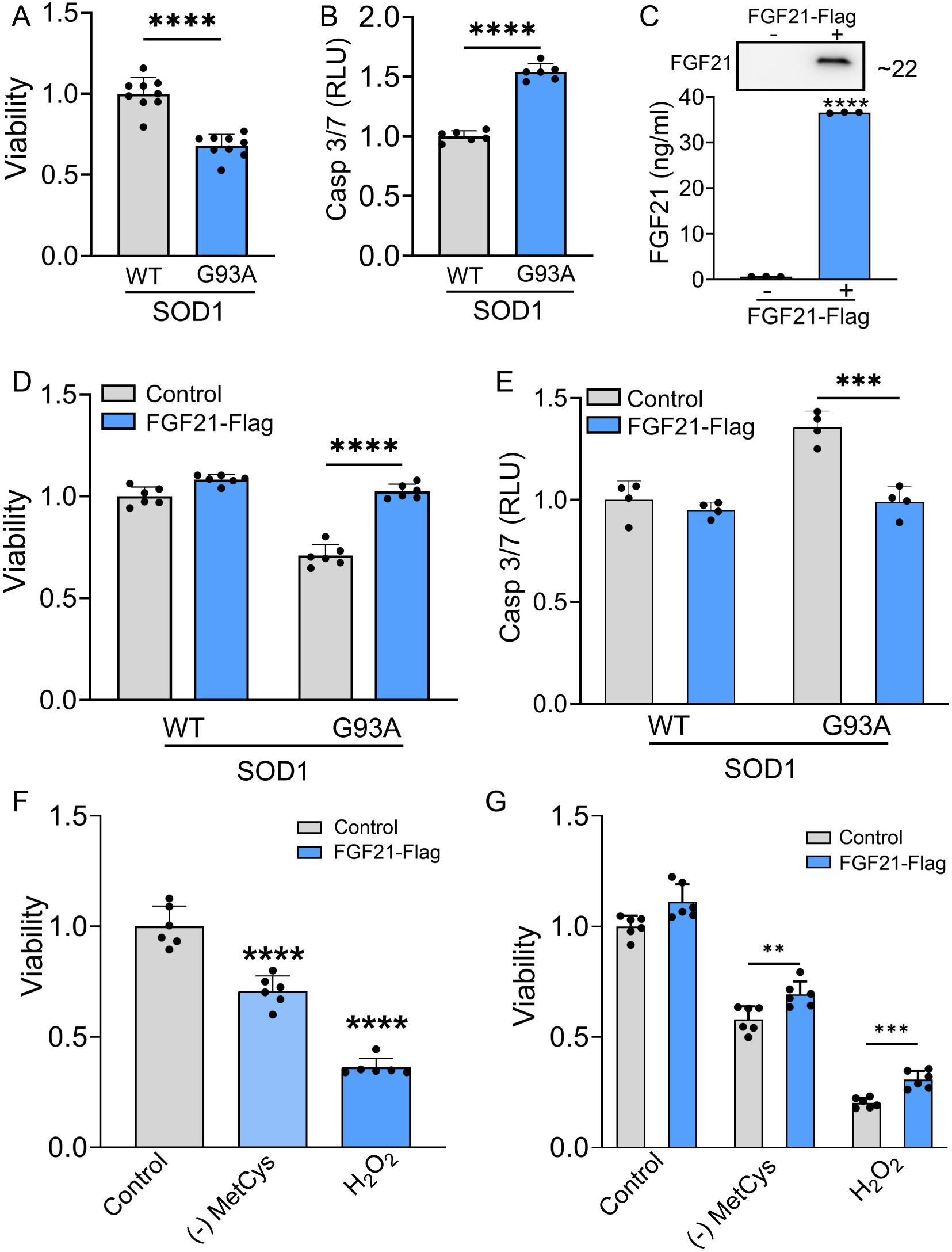
FGF21 mitigates cytotoxicity in NSC-34 motor neuron-like cells induced by SOD1^G93A^ or oxidative stress. **(A)** The viability of NSC-34 cells expressing FLAG-tagged WT-SOD1 or SOD1^G93A^ was determined using the Vialight assay. Viability for WT SOD1-transfected cells was set at 1. \*\*\*\**P <* 0.0001, unpaired two-tailed t-test. **(B)** Caspase activation was measured in the same cells and values were normalized to activity in WT SOD1-transfected cells which was set at 1. \*\*\*\**P <* 0.0001, unpaired two-tailed t-test. **(C)** FGF21 protein in the conditioned media of NSC-34 cells transfected with a FLAG-tagged FGF21 plasmid was detected by western blot (upper panel) and by ELISA (graph). \*\*\*\**P <* 0.0001; unpaired two-tailed t-test. **(D) and (E)** NSC-34 cells expressing either WT-SOD1 or SOD1^G93A^ were transfected with FLAG-FGF21 and assessed for viability as in (A) and Caspase-3/7 as in (B). \*\*\*\**P <* 0.0001, unpaired two-tailed t-test. **(F)** Cell viability was assessed in NSC-34 cells exposed to methionine-cystine (MetCys)-deprived media or treated with 100μM H_2_O_2_. \*\*\*\**P <* 0.0001; one-way ANOVA followed by Tukey’s multiple comparisons test. **(G)** NSC-34 cells transfected with FLAG-FGF21 (or empty vector) were subjected to stressors as described in (F) for 24h and then assayed for viability. \*\**P =* 0.007, \*\*\**P =* 0.0002; unpaired two-tailed t-test. Data points represent biological replicates and bars are the mean ± SD.

### FGF21 mitigates stress-induced toxicity in C2C12 muscle cells and promotes myogenesis

Since FGF21 is known to exert autocrine effects,^19^ and we found that atrophic myofibers in ALS muscle express FGF21 (Fig. 2), we next sought to assess the impact of FGF21 on muscle cells. Prior investigations have found that selective transgenic expression of SOD1^G93A^ in muscle induces oxidative stress leading to myofiber atrophy.^35^ We transfected C2C12 cells with SOD1^G93A^ and found that FGF21 expression was significantly reduced compared to WT SOD1, with the CM showing a ∼40% reduction in protein (Fig. 7A and B). *KLB* mRNA was not detected. The pattern of FGF21 suppression resembled ALS motor neurons (Fig. 5). With SOD1^G93A^ expression, there was a resultant loss of cell viability and an increase in Caspase-3/7 activity, each by 25%, compared to control (Fig. 7C; *P <* 0.0001). We transfected the muscle cells with Flag-tagged FGF21 which resulted in a marked increase in protein expression detected in the CM (Fig. 7D) and an attenuated loss of cell viability and apoptotic activity (Figs. 7E and 7F). We next assessed the effect of oxidative stress and found a significant induction of *FGF21* with both MetCys deprivation at 3.8-fold and H_2_O_2_ at 2.3-fold (Fig. 7G; *P <* 0.0001). *ATF4* and *PGC-1α* were also induced with these stressors by 1.5 to 2.5-fold (Supplementary Fig. 3C and 3D). Cell viability with MetCys deprivation led to a 75% reduction in viability and exposure to H_2_O_2_ resulted in a 50% reduction (Fig. 7H). Ectopic FGF21 expression improved viability by 100% for MetCys deprivation and 73% for H_2_O_2_ (Fig. 7I; *P <* 0.0001). Taken together, FGF21 is induced in C2C12 muscle cells expressing SOD1^G93A^ or when exposed to oxidative stress, resulting in cytotoxicity which can be reversed by FGF21.

**Figure 7.**
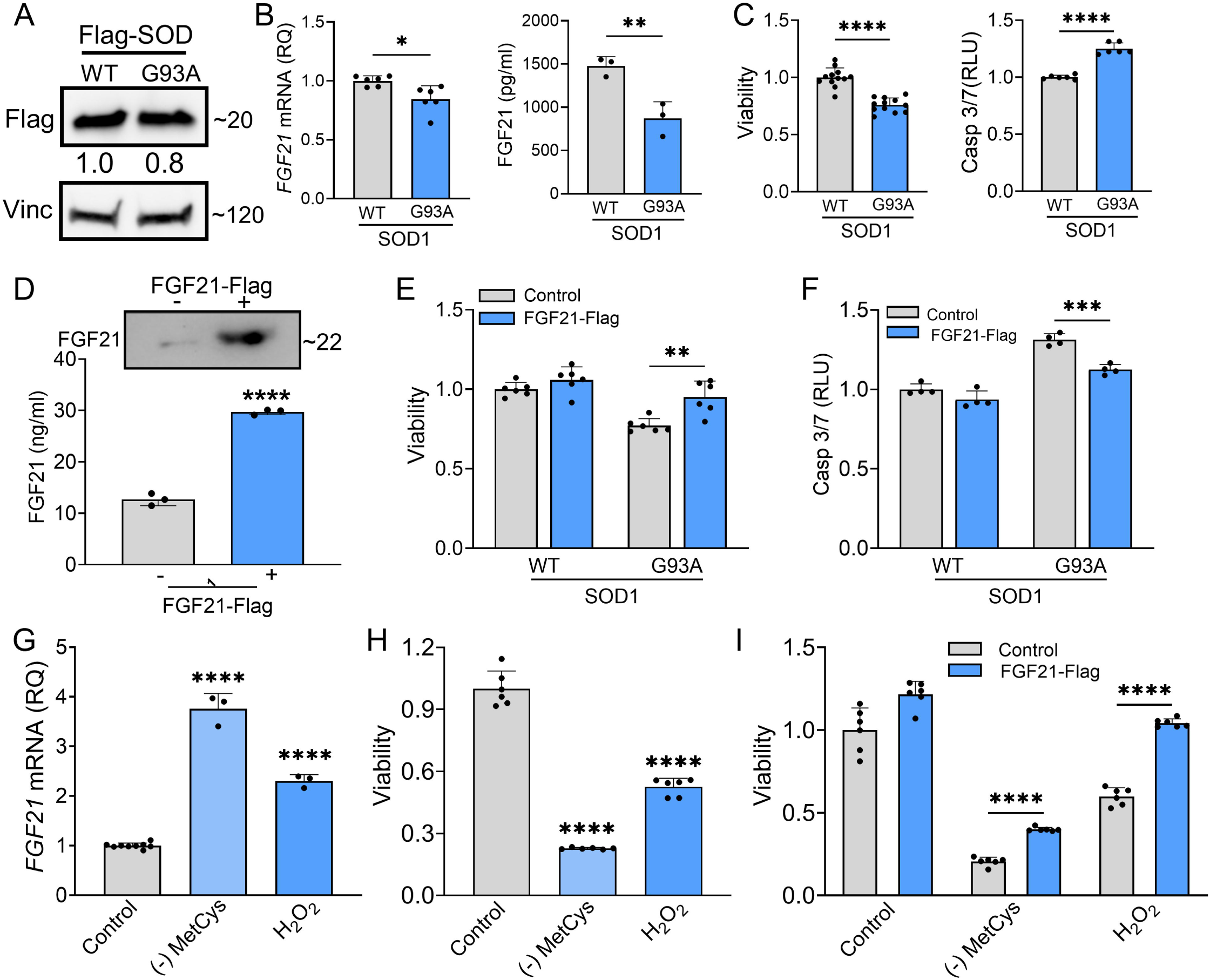
FGF21 mitigates cytotoxicity in C2C12 myoblasts induced by SOD1^G93A^ or oxidative stress. **(A)** C2C12 myoblasts were transfected with FLAG-tagged WT and SOD1^G93A^ expression plasmids and lysates were assessed by western blot using the antibodies indicated. Bands were quantitated by densitometry and a ratio to the loading control, vinculin, was calculated (shown between the two blots). **(B)** *FGF21* mRNA levels were measured in the same lysates (left panel) and FGF21 protein in the conditioned media was quantified by ELISA (right panel). \**P =* 0.011, \*\**P =* 0.008; unpaired two-tailed t-test. Secretory FGF21 from the conditioned media was quantified using ELISA (Right panel). (**C)** Viability of NSC-34 cells expressing FLAG-tagged WT-SOD1 or SOD1^G93A^ was determined using the Vialight assay (Left panel). Caspase activation was measured in the same cells and values were normalized to activity in WT SOD1-transfected cells which was set at 1 (right panel). \*\*\*\**P <* 0.0001, unpaired two-tailed t-test. **(D)** FGF21 protein in the conditioned media of C2C12 cells transfected with a FLAG-tagged FGF21 was detected by western blot (upper panel) and by ELISA (graph). Estimated size of the band (kDa) is shown to the right of the blot. \*\*\*\**P <* 0.0001; unpaired two-tailed t-test. **(E)** and **(F)** C2C12 myoblasts cells expressing either WT-SOD1 or SOD1^G93A^ were transfected with FGF21-FLAG and assessed for viability and Caspase-3/7 activity as in (C). \*\**P =* 0.003, \*\*\**P =* 0.0003; unpaired two-tailed t-test. **(G)** *FGF21* mRNA levels were quantified in C2C12 cells exposed to MetCys-deprived media or treated with 100μM H_2_O_2_ for 24 h. \*\*\*\**P <* 0.0001; one-way ANOVA followed by Tukey’s multiple comparisons test. **(H)** Cell viability was assessed in C2C12 myoblasts exposed to stressors as described in (G). \*\*\*\**P <* 0.0001; one-way ANOVA followed by Tukey’s multiple comparisons test. **(I)** C2C12 cells transfected with FGF21-FLAG (or empty vector) were subjected to stressors as described in (G) for 24h and then assayed for viability. \*\*\*\**P <* 0.0001; unpaired two-tailed t-test. Data points represent biological replicates and bars are the mean ± SD.

We further investigated the impact of FGF21 on myogenesis as prior studies in preclinical models of ALS have linked myogenesis to slower disease progression.^36,37^ We induced myogenic differentiation in C2C12 cells over 96 h as reflected by an increased fusion index (Fig. 8A) and induction of MHC (Fig. 8B). Over the same time interval, there was a progressive and significant increase in both *FGF21* mRNA within cells and secreted FGF21 in the CM (Fig. 8C). When FGF21 was ectopically expressed in C2C12 cells, there was a significant increase in myogenic differentiation (Fig. 8D) as indicated by a significantly higher fusion index (Fig. 8E; *P =* 0.008) and a 2.2-fold induction of MHC (Fig. 8F). Cell proliferation was then assessed using an MTS assay (Fig. 8G). Just prior to seeding, FGF21 was quantitated in the CM by ELISA and showed a ∼7-fold increase in FGF21 in the FGF21-FLAG transfected cells versus empty vector control. We observed a significant increase in proliferation in FGF21-FLAG transfected cells at all time intervals tested, reaching 25% at 72 h (*P <* 0.0001). In summary, FGF21 induction occurs as part of the myogenic differentiation program, where it appears to play a positive role in myogenesis.

**Figure 8.**
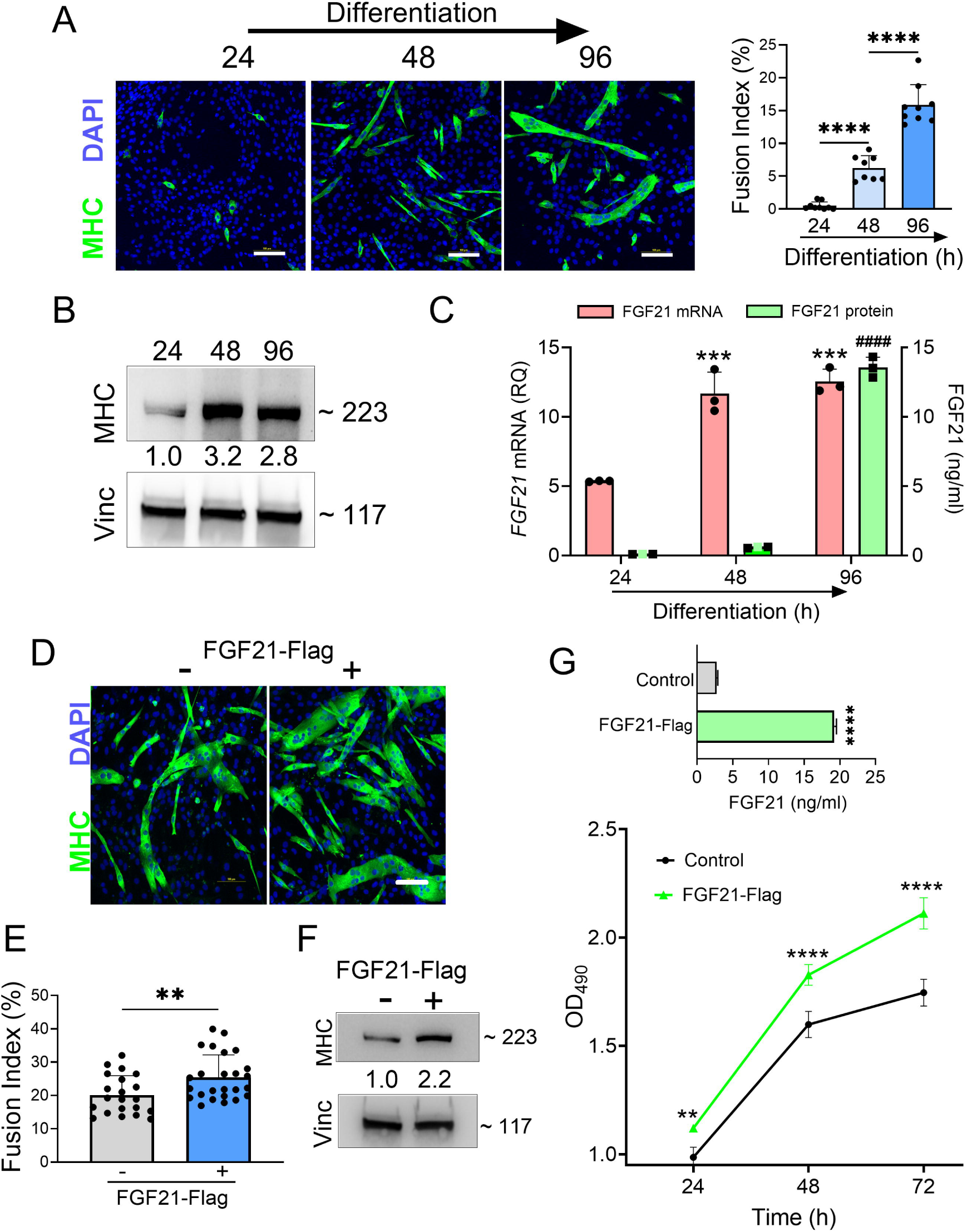
FGF21 is upregulated during myogenesis and facilitates myogenic differentiation of C2C12 myoblasts. **(A)** C2C12 myoblasts were treated with DM for various time intervals and immunostained with an anti-MHC antibody followed by DAPI counterstaining. Myotube formation was detected by MHC-positive staining. Scale bars, 100 µM. The fusion index (%) was quantified as described in the methods (right panel). **(B)** C2C12 myoblasts treated with DM were lysed at specific time intervals and immunoblotted with antibodies against MHC and vinculin. Densitometry values (24 h time interval was set at 1) are shown. **(C)** *FGF21* mRNA levels were quantified from the lysates and FGF21 protein from conditioned media. \*\*\**P =* 0.0005 comparing to the 24 h time interval, ^####^*P <* 0.0001 comparing to the 24 and 48 h time intervals; one-way ANOVA followed by Tukey’s multiple comparisons test. **(D)** C2C12 myoblasts were transfected with an FGF21-FLAG plasmid and cultured in DM for 96 h. Myotube formation was assessed by MHC-positive staining as in (A). Scale bar, 100 µM. **(E)** Fusion index for transfected C2C12 cells. \*\**P =* 0.008; unpaired two-tailed t-test. **(F)** Immunoblot analysis for MHC and vinculin was performed as described in (B). **(G)** FGF21 levels in the conditioned media from C2C12 myoblasts transfected with FGF21-FLAG or empty vector control were quantified by ELISA (upper graph). \*\*\*\**P* < 0.0001; unpaired two-tailed t test. Cells were reseeded and cultured in growth medium (GM) for 72 h. Cell proliferation was assessed at indicated time intervals using MTS (lower graph). \*\**P* = 0.007, \*\*\*\**P* < 0.0001; one-way ANOVA followed by Tukey’s multiple comparisons test. Data points represent biological replicates and bars are the mean ± SD.

## Discussion

In this study, we found that FGF21 is prominently upregulated in atrophic myofibers in patients with ALS and increases with disease progression. FGF21 is also increased in plasma of ALS patients as a group, but disproportionately in those with slower clinical progression and higher BMI. We show that the RNA expression of the key co-receptor, β-Klotho, is upregulated in ALS motor neurons at baseline and that both *FGF21* and *KLB* are induced by oxidative stress with FGF21 appearing to provide a protective effect. In muscle cells, FGF21 promotes myogenesis and exerts a similar protective effect against cytotoxic stress. Taken together, these findings identify FGF21 as a biomarker that provides insight into the clinical heterogeneity and pathophysiology of ALS while at the same time identifying it as a potential target for future therapeutic development.

Although liver is the predominant source of FGF21 in physiological states, mitochondrial dysfunction, as seen in skeletal muscle at the earliest stages of ALS, is a potent stimulus for FGF21 induction.^16,38,39^ In mitochondrial myopathies, elevated serum FGF21 from skeletal muscle is now recognized as a diagnostic biomarker.^40–42^ Our data suggest that one source of elevated FGF21 in plasma is from skeletal muscle as the protein levels were high (more than 2-fold higher than plasma levels and 20-fold higher than spinal cord tissue) and immunohistochemistry showed substantial extravasation into the endomysial space (Figs. 1-3). Unlike most FGF family members, FGF21 has a lower affinity for heparan sulfate glycosaminoglycans^33^ which are present in the basal lamina surrounding muscle fibers,^43^ and thus less likely to get sequestered. This is in contrast to FGF23, a related family member that we previously identified in ALS muscle samples but could not detect in plasma samples.^21^ Under physiological conditions, liver is the predominant source of circulating FGF21.^20,33^ Our findings in the SOD1^G93A^ mouse, however, suggest that muscle is a major source as there was a ∼2-fold higher expression level versus liver (Supplementary Fig. 2). Interestingly, we found increased FGF21 protein in spinal cord, but no overall increase in *FGF21* mRNA, raising the possibility that FGF21 may be coming from the periphery. FGF21 can readily cross the blood-brain barrier, and elevated plasma levels have previously been shown to correlate with elevated cerebrospinal fluid (CSF) levels in humans.^44,45^ On the other hand, we detected increase *FGF21* transcripts in whole spinal cord of the SOD1^G93A^ mouse at a very early, presymptomatic, stage (∼ 6 weeks; Fig. 1). Others have detected induction of *FGF21* in a subset motor neurons in the SOD1^G93A^ and TDP43 mouse models in late presymptomatic/early symptomatic stages (11-12 weeks post natal).^46,47^ Although protein levels were not measured, one of the studies found a high association of *FGF21* with ribosomes in motor neurons.^47^ It is possible that post-transcriptional/translational mechanisms are at play within the CNS, particularly since the protein half-life of FGF21 is short.^33^

The potential for FGF21 to track disease progression is supported by several lines of evidence. First, since a major hallmark of ALS is progressive muscle atrophy,^30^ FGF21 would be expected to increase over time because of its correlation with atrophied myofibers (Fig. 2) and HDAC4, a muscle biomarker that associates with progressive muscle denervation in ALS.^28^ Indeed, this is supported by the nearly 8-fold increase in muscle *FGF21* mRNA levels in end-stage ALS muscle versus muscle biopsy samples at earlier stages (Fig. 1). There was also a progressive increase in *FGF21* mRNA in ALS mouse muscle starting in the presymptomatic stages (Fig. 1). If muscle FGF21 were the only source for increased plasma FGF21, however, higher levels might be expected to associate with more advanced disease (i.e. more muscle atrophy) as has been observed with mitochondrial myopathies.^41,48^ In our longitudinal study of 16 patients, however, those with high FGF21 plasma levels showed slower disease progression and longer survival (Fig. 3). This raises the possibility that in some ALS patients, other sources contribute to circulating FGF21 such as liver. A notable feature of the subset with high circulating FGF21 levels was a higher mean BMI which verged on obesity. Prior studies have shown a strong positive correlation between serum FGF21 levels and BMI, particularly increased fat mass, with liver being the likely source.^44,49^ Increased BMI has consistently been linked to longer survival in ALS in a number of clinical studies.^50–53^ In the Pooled Resource Open-Access ALS Clinical Trials database (PRO-ACT), patients with a BMI between 25 – 30 kg/M^2^ (similar to the mean value of 29.2 kg/M^2^ in the high FGF21 subset reported here) had a 35% reduced risk of dying compared to patients with BMI < 25 kg/M^2^ (the low FGF21 group in our study had a mean BMI of 22.6 kg/M^2^).^52^ In a pilot study of ALS patients, we previously found that low percentage body fat and loss of total fat mass correlated with faster monthly declines in ALSFRS-R scores.^54^ In another study, patients with low visceral fat mass at baseline had significantly faster disease progression.^55^ Interestingly, the majority of patients in our study with low circulating FGF21 levels had bulbar onset ALS (Supplementary Table 4) which is associated with shorter lifespan and weight loss prior to diagnosis (most likely due to decreased nutritional intake).^56,57^

A limitation of our plasma FGF21 study is the small number of patients, which in a heterogeneous disorder such as ALS, can lead to premature conclusions. A larger ALS cohort will be required for validation. The findings are timely, however, given the increasing attention to dysregulation of energy metabolism as a key driver of ALS disease progression.^6,58,59^ FGF21 is a master regulator of metabolic and nutrient homeostasis and a key hormone in adapting to states of energy deprivation.^33^ While it is well established that FGF21 is a key regulator of fatty acid oxidation in the liver,^19^ less is known about this function in skeletal muscle or in the CNS.^20^ Preclinical studies in the ALS mouse suggest that fatty acid oxidation becomes a major source of energy in skeletal muscle in the SOD1^G93A^.^60–62^ In myotubes derived from human ALS muscle, increased fatty acid oxidation capacity was associated with slower disease progression.^61^ It is interesting to note that a recent study on macronutrients in ALS suggests that a high glycemic index diet, which is a potent stimulus for FGF21 induction by the liver,^63^ is linked to slower disease progression in patients with ALS.^64^

The biology of FGF21 is complex because of its broad and divergent effects on different organs that vary depending on the physiological or pathological context.^19,33^ In our study, we found increased FGF21 in skeletal muscle, spinal cord and in the circulation, suggesting that multiple mechanisms may impact ALS pathophysiology, including stress response, autophagy, energy metabolism, inflammation, and trophic effects. Because of the increase in FGF21 in the CNS (Fig. 1) and the previously reported correlation between higher CSF FGF21 levels and high serum FGF21 (and concomitant BMI)^44^, potential target cell populations affected in ALS are also broad. This impact is underscored by preclinical studies in Parkinson’s, Alzheimer’s and other CNS disease where FGF21 promotes neuroprotection and mitigates neurodegeneration through effects on different cell populations which crossover to ALS. This includes suppression of glial activation and neuroinflammation, glutamate excitotoxicity, reversal of defective astrocyte-neuron lactate shuttling, increased neuronal survival, and enhanced myelin regeneration mediated by oligodendrocytes.^65–71^ We focused on the potential trophic effects of FGF21 based on prior studies showing a direct protective effect of FGF family members, FGF1 and 2, on motor neurons after spinal cord injury.^72,73^ A possible benefit of FGF21 signaling was also suggested in a prior study with R1Mab1, an FGFR1-targeting antibody that activates the receptor, in the SOD1^G93A^ mouse where there was mild amelioration of motor phenotype.^74^

A key component of FGF21 signaling is the co-receptor, β-klotho (KLB), which exerts cellular specificity of signaling as the FGF receptors are nearly ubiquitous.^33^ Our detection of *KLB* in iPSC-derived human motor neurons (Fig. 5) indicates a high likelihood that FGF21 signaling is relevant to motor neuron physiology. Evidence for FGF21 signaling as a stress response in motor neurons was further supported by our finding of *KLB* and *FGF21* induction in NSC-34 cells with oxidative stress, either by exposure to H_2_O_2_ or deprivation of MetCys (Fig. 5). Interestingly, we found more than a 3-fold increase in *KLB* mRNA in ALS motor neurons and this pattern was recapitulated in NSC-34 motor neuron-like cells expressing SOD1^G93A^ (Fig. 5). This baseline increase suggests that the trigger for FGF signaling is already present in ALS motor neurons at the time of differentiation. Oxidative stress has previously been observed in differentiated motor neurons from ALS patients using intracellular biosensors, and this might reflect an early and intrinsic vulnerability of ALS motor neurons.^75,76^ At the same time, however, basal *FGF21* levels were reduced suggesting an impairment in the FGF21-KLB axis. In ALS tissue samples, we observed an upregulation of *KLB* in muscle biopsies but a downregulation in end-stage muscle and spinal cord (Fig. 4). This pattern could be consistent with disease-associated loss of motor neurons. More recently, KLB was identified in muscle cells, albeit at low levels,^20^ so our finding of reduced *KLB* in end-stage muscle could also be related to loss of myofibers. Nonetheless, the concomitant higher FGF21 levels in ALS muscle and spinal cord at end-stage suggest an ongoing adaptive response to disease progression. A trophic effect of FGF21 on SOD1^G93A^ motor neurons was supported by the reversal of cytotoxicity and apoptosis with ectopic expression (Fig. 6). Potential mechanisms for this rescue effect include attenuation of oxidative stress, modulation of autophagy, improved energy metabolism and mitochondrial function.^66,67^

Selective transgenic expression of SOD1^G93A^ in skeletal muscle has been shown to promote mitochondrial dysfunction, oxidative stress, and progressive atrophy.^35^ We found a similar toxic effect in muscle cells and that ectopic FGF21 expression reversed this toxicity (Fig. 7). FGF21 activates mTOR signaling in muscle cells leading to improved mitochondrial function, energy metabolism and reversal of mitochondrial impairment by oxidative stress.^77^ In addition to a rescue effect, we also found that FGF21 promoted myogenesis, including enhanced proliferation and differentiation of C2C12 myoblasts (Fig. 8), as observed by other investigators.^78,79^ A prior study assessing satellite cells from human ALS muscle tissue revealed defects in myogenesis, particularly differentiation and maturation.^80^ Defects in myogenesis were also observed in satellite cells derived from SOD1^G93A^ mice at early pre-symptomatic stages^81^ and in C2C12 cells expressing SOD1^G93A^.^82^ Recently, increased myogenesis was identified as a key mechanism for the slow-progressing phenotype in SOD1^G93A^ mice from the C57 background.^36,37^ One study found that that FGF21 drives the molecular transition of myoblasts to an aerobic (slow twitch fiber-like) phenotype,^78^ a process that occurs in ALS muscle due to the vulnerability and early degeneration of motor neurons innervating fast-twitch (Type II) muscle fibers.^83,84^

FGF21 has been identified as a biomarker of frailty because of its association with age-related metabolic disorders.^85–87^ Aging is a key risk factor for ALS, and many pathophysiological features of ALS overlap with normal aging patterns including mitochondrial dysfunction, oxidative stress and inflammation.^88^ The role of FGF21 signaling in these age-associated processes is unclear. Many studies support a mitigating effect of FGF21 on aging, mitochondrial function, muscle atrophy and function via its effects on metabolism.^85,86^ Other studies suggest FGF21 signaling exerts a negative effect on myogenesis and promotes atrophy in different physiological and disease contexts including fasting, aging, intrauterine growth restriction, and short-term pharmacological administration.^89–92^ These conflicting findings underscore the complexity of FGF21 biology and the need for further investigation, including the role of β-klotho.^87^

In summary, we have identified FGF21 as a novel biomarker in ALS that is detected in multiple compartments including muscle, spinal cord, and the circulation. It is strongly expressed in atrophied myofibers, and high plasma levels associate with slower disease progression. For the first time, we show that the FGF21-KLB axis is a relevant stress response pathway in motor neurons and that it mitigates cytotoxicity induced by oxidative stress and co-expression of mutant SOD1. It has a similar protective effect in muscle cells where it also promotes myogenesis. Future studies will need to test FGF21 using *in vivo* models of ALS, including tissue-specific or systemic delivery, to validate the mitigating effects it appears to have *in cellulo* and to assess its potential as a novel therapy in ALS. In the clinic, plasma FGF21 levels might have value as a prognostic biomarker but will need validation in larger multi-center studies.

## Author Contributions

A.G. and P.H.K. contributed to the conception and design of the study. PHK, YS, MK, and NJ developed the ALS and control tissue sample repository at UAB. A.G, Y.S., R.S., M.K., N.J., B.K.S., R.H., J.D.T.D.S.P., J.S.S., M.A., E.H.V., and P.H.K. contributed to data acquisition. A.G., Y.S., K.S., A.T.M., S.A.A., and P.H.K. contributed to data analysis and interpretation; A.G., and P.H.K. contributed to drafting the text and preparing the figures.

## Acknowledgements

Some of the ALS tissue specimens were provided by the Department of Veterans Affairs Biorepository (VA Merit review BX002466). Normal control tissues were provided by the Tissue Procurement Shard Resource and the UAB Tissue Biorepository. We are also grateful to our patients who courageously agreed to donate tissue samples post-mortem to advance ALS research.

## Ethical Statement

Collection of all human tissue and blood samples was approved by the following Institutional Review Boards: UAB (091222037 and X100908007), Birmingham VA Medical Center (151217001) and Cedars-Sinai (21505).

## Consent

Written informed consent was obtained from patients who participated in these studies. For post-mortem tissue collection, informed consent was also obtained from the next-of-kin at the time of death.

## References

1. Kazamel M, Cutter G, Claussen G, et al. Epidemiological features of amyotrophic lateral sclerosis in a large clinic-based African American population. Amyotroph Lateral Scler Frontotemporal Degener. Sep 2013;14(5-6):334–7. doi:10.3109/21678421.2013.770030

2. Hardiman O, van den Berg LH, Kiernan MC. Clinical diagnosis and management of amyotrophic lateral sclerosis. 10.1038/nrneurol.2011.153. Nat Rev Neurol. Oct 11 2011;7(11):639–49. doi:10.1038/nrneurol.2011.153

3. Verma S, Khurana S, Vats A, et al. Neuromuscular Junction Dysfunction in Amyotrophic Lateral Sclerosis. Mol Neurobiol. Mar 2022;59(3):1502–1527. doi:10.1007/s12035-021-02658-6

4. King PH. Skeletal muscle as a molecular and cellular biomarker of disease progression in amyotrophic lateral sclerosis: a narrative review. Neural Regen Res. Apr 2024;19(4):747–753. doi:10.4103/1673-5374.382226

5. Moloney EB, de Winter F, Verhaagen J. ALS as a distal axonopathy: molecular mechanisms affecting neuromuscular junction stability in the presymptomatic stages of the disease. Front Neurosci. 2014;8:252. doi:10.3389/fnins.2014.00252

6. Nelson AT, Trotti D. Altered Bioenergetics and Metabolic Homeostasis in Amyotrophic Lateral Sclerosis. Neurotherapeutics. 2022/06/30 2022;doi:10.1007/s13311-022-01262-3

7. Kwan T, Kazamel M, Thoenes K, Si Y, Jiang N, King PH. Wnt antagonist FRZB is a muscle biomarker of denervation atrophy in amyotrophic lateral sclerosis. Sci Rep. Oct 7 2020;10(1):16679. doi:10.1038/s41598-020-73845-z

8. Si Y, Cui X, Kim S, et al. Smads as muscle biomarkers in amyotrophic lateral sclerosis. Ann Clin Transl Neurol. Oct 2014;1(10):778–87. doi:10.1002/acn3.117

9. Si Y, Kazamel M, Kwon Y, et al. The vitamin D activator CYP27B1 is upregulated in muscle fibers in denervating disease and can track progression in amyotrophic lateral sclerosis. J Steroid Biochem Mol Biol. Jun 2020;200:105650. doi:10.1016/j.jsbmb.2020.105650

10. Si Y, Kim S, Cui X, et al. Transforming Growth Factor Beta (TGF-beta) Is a Muscle Biomarker of Disease Progression in ALS and Correlates with Smad Expression. PLoS One. 2015;10(9):e0138425. doi:10.1371/journal.pone.0138425

11. Tsitsipatis D, Mazan-Mamczarz K, Si Y, et al. Transcriptomic analysis of human ALS skeletal muscle reveals a disease-specific pattern of dysregulated circRNAs. Aging. Dec 30 2022;14(24):9832–9859. doi:10.18632/aging.204450

12. Si Y, Cui X, Crossman DK, et al. Muscle microRNA signatures as biomarkers of disease progression in amyotrophic lateral sclerosis. Neurobiol Dis. Jun 2018;114:85–94. doi:10.1016/j.nbd.2018.02.009

13. Lopez MA, Si Y, Hu X, et al. Smad8 Is Increased in Duchenne Muscular Dystrophy and Suppresses miR-1, miR-133a, and miR-133b. Int J Mol Sci. Jul 7 2022;23(14):7515. doi:10.3390/ijms23147515

14. Darabid H, Perez-Gonzalez AP, Robitaille R. Neuromuscular synaptogenesis: coordinating partners with multiple functions. Nature Reviews Neuroscience. 2014/11/01 2014;15(11):703–718. doi:10.1038/nrn3821

15. Benatar M, Boylan K, Jeromin A, et al. ALS biomarkers for therapy development: State of the field and future directions. Muscle Nerve. Feb 2016;53(2):169–82. doi:10.1002/mus.24979

16. Tezze C, Romanello V, Sandri M. FGF21 as Modulator of Metabolism in Health and Disease. Review. Front Physiol. 2019-April-17 2019;10 doi:10.3389/fphys.2019.00419

17. Jena J, García-Peña LM, Pereira RO. The roles of FGF21 and GDF15 in mediating the mitochondrial integrated stress response. Front Endocrinol (Lausanne). 2023;14:1264530. doi:10.3389/fendo.2023.1264530

18. Yan B, Mei Z, Tang Y, et al. FGF21-FGFR1 controls mitochondrial homeostasis in cardiomyocytes by modulating the degradation of OPA1. Cell Death Dis. May 8 2023;14(5):311. doi:10.1038/s41419-023-05842-9

19. Fisher FM, Maratos-Flier E. Understanding the Physiology of FGF21. Annu Rev Physiol. 2016;78:223–41. doi:10.1146/annurev-physiol-021115-105339

20. Sun H, Sherrier M, Li H. Skeletal Muscle and Bone - Emerging Targets of Fibroblast Growth Factor-21. Frontiers in physiology. 2021;12:625287. doi:10.3389/fphys.2021.625287

21. Si Y, Kazamel M, Benatar M, et al. FGF23, a novel muscle biomarker detected in the early stages of ALS. Sci Rep. Jun 8 2021;11(1):12062. doi:10.1038/s41598-021-91496-6

22. Lu L, Zheng L, Viera L, et al. Mutant Cu/Zn-superoxide dismutase associated with amyotrophic lateral sclerosis destabilizes vascular endothelial growth factor mRNA and downregulates its expression. J Neurosci. Jul 25 2007;27(30):7929–38. doi:10.1523/JNEUROSCI.1877-07.2007

23. Selvaraj BT, Livesey MR, Zhao C, et al. C9ORF72 repeat expansion causes vulnerability of motor neurons to Ca2+-permeable AMPA receptor-mediated excitotoxicity. Nature Communications. 2018/01/24 2018;9(1):347. doi:10.1038/s41467-017-02729-0

24. Yang S, Wijegunawardana D, Sheth U, et al. Aberrant splicing exonizes C9ORF72 repeat expansion in ALS/FTD. bioRxiv. Nov 14 2023;doi:10.1101/2023.11.13.566896

25. Lu L, Wang S, Zheng L, et al. Amyotrophic lateral sclerosis-linked mutant SOD1 sequesters Hu antigen R (HuR) and TIA-1-related protein (TIAR): implications for impaired post-transcriptional regulation of vascular endothelial growth factor. J Biol Chem. Dec 4 2009;284(49):33989–98. doi:10.1074/jbc.M109.067918

26. Noë S, Corvelyn M, Willems S, et al. The Myotube Analyzer: how to assess myogenic features in muscle stem cells. Skelet Muscle. Jun 10 2022;12(1):12. doi:10.1186/s13395-022-00297-6

27. Logroscino G, Traynor BJ, Hardiman O, et al. Incidence of amyotrophic lateral sclerosis in Europe. J Neurol Neurosurg Psychiatry. April 1, 2010 2010;81(4):385–390. doi:10.1136/jnnp.2009.183525

28. Bruneteau G, Simonet T, Bauche S, et al. Muscle histone deacetylase 4 upregulation in amyotrophic lateral sclerosis: potential role in reinnervation ability and disease progression. Brain. Aug 2013;136(Pt 8):2359–68. doi:10.1093/brain/awt164

29. Vinsant S, Mansfield C, Jimenez-Moreno R, et al. Characterization of early pathogenesis in the SOD1(G93A) mouse model of ALS: part II, results and discussion. Brain Behav. Jul 2013;3(4):431–57. doi:10.1002/brb3.142

30. Al-Sarraj S, King A, Cleveland M, et al. Mitochondrial abnormalities and low grade inflammation are present in the skeletal muscle of a minority of patients with amyotrophic lateral sclerosis; an observational myopathology study. journal article. Acta Neuropathol Commun. Dec 14 2014;2(1):165. doi:10.1186/s40478-014-0165-z

31. Post A, Dam WA, Sokooti S, et al. Circulating FGF21 Concentration, Fasting Plasma Glucose, and the Risk of Type 2 Diabetes: Results From the PREVEND Study. The Journal of Clinical Endocrinology & Metabolism. 2022;108(6):1387–1393. doi:10.1210/clinem/dgac729

32. Labra J, Menon P, Byth K, Morrison S, Vucic S. Rate of disease progression: a prognostic biomarker in ALS. J Neurol Neurosurg Psychiatry. Jun 2016;87(6):628–32. doi:10.1136/jnnp-2015-310998

33. BonDurant LD, Potthoff MJ. Fibroblast Growth Factor 21: A Versatile Regulator of Metabolic Homeostasis. Annu Rev Nutr. 2018;38(Volume 38, 2018):173–196. 10.1146/annurev-nutr-071816-064800

34. Gille JJP, Joenje H. Cell culture models for oxidative stress: superoxide and hydrogen peroxide versus normobaric hyperoxia. Mutation Research/DNAging. 1992/09/01/ 1992;275(3):405–414. 10.1016/0921-8734(92)90043-O

35. Dobrowolny G, Aucello M, Rizzuto E, et al. Skeletal muscle is a primary target of SOD1G93A-mediated toxicity. Cell metabolism. Nov 2008;8(5):425–36. doi:10.1016/j.cmet.2008.09.002

36. Margotta C, Fabbrizio P, Ceccanti M, et al. Immune-mediated myogenesis and acetylcholine receptor clustering promote a slow disease progression in ALS mouse models. Inflamm Regen. Mar 9 2023;43(1):19. doi:10.1186/s41232-023-00270-w

37. Trolese MC, Scarpa C, Melfi V, et al. Boosting the peripheral immune response in the skeletal muscles improved motor function in ALS transgenic mice. Mol Ther. Aug 3 2022;30(8):2760–2784. doi:10.1016/j.ymthe.2022.04.018

38. Loeffler JP, Picchiarelli G, Dupuis L, Gonzalez De Aguilar JL. The Role of Skeletal Muscle in Amyotrophic Lateral Sclerosis. Brain Pathol. Mar 2016;26(2):227–36. doi:10.1111/bpa.12350

39. Zhou J, Yi J, Fu R, et al. Hyperactive intracellular calcium signaling associated with localized mitochondrial defects in skeletal muscle of an animal model of amyotrophic lateral sclerosis. J Biol Chem. Jan 1 2010;285(1):705–12. doi:10.1074/jbc.M109.041319

40. Suomalainen A, Elo JM, Pietiläinen KH, et al. FGF-21 as a biomarker for muscle-manifesting mitochondrial respiratory chain deficiencies: a diagnostic study. Lancet Neurol. Sep 2011;10(9):806–18. doi:10.1016/s1474-4422(11)70155-7

41. Lehtonen JM, Forsström S, Bottani E, et al. FGF21 is a biomarker for mitochondrial translation and mtDNA maintenance disorders. Neurology. Nov 29 2016;87(22):2290–2299. doi:10.1212/wnl.0000000000003374

42. Tyynismaa H, Carroll CJ, Raimundo N, et al. Mitochondrial myopathy induces a starvation-like response. Human Molecular Genetics. 2010;19(20):3948–3958. doi:10.1093/hmg/ddq310

43. Jenniskens GJ, Oosterhof A, Brandwijk R, Veerkamp JH, van Kuppevelt TH. Heparan sulfate heterogeneity in skeletal muscle basal lamina: demonstration by phage display-derived antibodies. J Neurosci. Jun 1 2000;20(11):4099–111. doi:10.1523/JNEUROSCI.20-11-04099.2000

44. Tan BK, Hallschmid M, Adya R, Kern W, Lehnert H, Randeva HS. Fibroblast Growth Factor 21 (FGF21) in Human Cerebrospinal Fluid: Relationship With Plasma FGF21 and Body Adiposity. Diabetes. 2011;60(11):2758–2762. doi:10.2337/db11-0672

45. Hsuchou H, Pan W, Kastin AJ. The fasting polypeptide FGF21 can enter brain from blood. Peptides. 2007/12/01/ 2007;28(12):2382–2386. 10.1016/j.peptides.2007.10.007

46. Henriques A, Kastner S, Chatzikonstantinou E, et al. Gene expression changes in spinal motoneurons of the SOD1(G93A) transgenic model for ALS after treatment with G-CSF. Front Cell Neurosci. 2014;8:464. doi:10.3389/fncel.2014.00464

47. Shadrach JL, Stansberry WM, Milen AM, et al. Translatomic analysis of regenerating and degenerating spinal motor neurons in injury and ALS. iScience. Jul 23 2021;24(7):102700. doi:10.1016/j.isci.2021.102700

48. Koene S, de Laat P, van Tienoven DH, et al. Serum FGF21 levels in adult m.3243A>G carriers: clinical implications. Neurology. Jul 8 2014;83(2):125–33. doi:10.1212/wnl.0000000000000578

49. Dushay J, Chui PC, Gopalakrishnan GS, et al. Increased Fibroblast Growth Factor 21 in Obesity and Nonalcoholic Fatty Liver Disease. Gastroenterology. 2010/08/01/ 2010;139(2):456–463. 10.1053/j.gastro.2010.04.054

50. Witzel S, Wagner M, Zhao C, et al. Fast versus slow disease progression in amyotrophic lateral sclerosis–clinical and genetic factors at the edges of the survival spectrum. Neurobiol Aging. 2022/11/01/ 2022;119:117–126. 10.1016/j.neurobiolaging.2022.07.005

51. Dardiotis E, Siokas V, Sokratous M, et al. Body mass index and survival from amyotrophic lateral sclerosis: A meta-analysis. Neurol Clin Pract. Oct 2018;8(5):437–444. doi:10.1212/CPJ.0000000000000521

52. Atassi N, Berry J, Shui A, et al. The PRO-ACT database: design, initial analyses, and predictive features. Neurology. Nov 4 2014;83(19):1719–25. doi:10.1212/WNL.0000000000000951

53. Traxinger K, Kelly C, Johnson BA, Lyles RH, Glass JD. Prognosis and epidemiology of amyotrophic lateral sclerosis: Analysis of a clinic population, 1997-2011. Neurol Clin Pract. Aug 2013;3(4):313–320. doi:10.1212/CPJ.0b013e3182a1b8ab

54. Lee I, Kazamel M, McPherson T, et al. Fat mass loss correlates with faster disease progression in amyotrophic lateral sclerosis patients: Exploring the utility of dual-energy x-ray absorptiometry in a prospective study. PLoS One. 2021;16(5):e0251087. doi:10.1371/journal.pone.0251087

55. Li J-Y, Sun X-H, Cai Z-Y, et al. Correlation of weight and body composition with disease progression rate in patients with amyotrophic lateral sclerosis. Sci Rep. 2022/08/02 2022;12(1):13292. doi:10.1038/s41598-022-16229-9

56. Chiò A, Logroscino G, Hardiman O, et al. Prognostic factors in ALS: A critical review. Amyotroph Lateral Scler. Oct-Dec 2009;10(5-6):310–23. doi:10.3109/17482960802566824

57. Moglia C, Calvo A, Grassano M, et al. Early weight loss in amyotrophic lateral sclerosis: outcome relevance and clinical correlates in a population-based cohort. J Neurol Neurosurg Psychiatry. Jun 2019;90(6):666–673. doi:10.1136/jnnp-2018-319611

58. Jésus P, Fayemendy P, Nicol M, et al. Hypermetabolism is a deleterious prognostic factor in patients with amyotrophic lateral sclerosis. Eur J Neurol. Jan 2018;25(1):97–104. doi:10.1111/ene.13468

59. Steyn FJ, Ioannides ZA, van Eijk RPA, et al. Hypermetabolism in ALS is associated with greater functional decline and shorter survival. J Neurol Neurosurg Psychiatry. Oct 2018;89(10):1016–1023. doi:10.1136/jnnp-2017-317887

60. Dupuis L, Oudart H, Rene F, Gonzalez de Aguilar JL, Loeffler JP. Evidence for defective energy homeostasis in amyotrophic lateral sclerosis: benefit of a high-energy diet in a transgenic mouse model. Research Support, Non-U.S. Gov’t. Proc Natl Acad Sci USA. Jul 27 2004;101(30):11159–64. doi:10.1073/pnas.0402026101

61. Steyn FJ, Li R, Kirk SE, et al. Altered skeletal muscle glucose-fatty acid flux in amyotrophic lateral sclerosis. Brain Commun. 2020;2(2):fcaa154. doi:10.1093/braincomms/fcaa154

62. Palamiuc L, Schlagowski A, Ngo ST, et al. A metabolic switch toward lipid use in glycolytic muscle is an early pathologic event in a mouse model of amyotrophic lateral sclerosis. EMBO Mol Med. May 2015;7(5):526–46. doi:10.15252/emmm.201404433

63. Solon-Biet SM, Cogger VC, Pulpitel T, et al. Defining the Nutritional and Metabolic Context of FGF21 Using the Geometric Framework. Cell metabolism. Oct 11 2016;24(4):555–565. doi:10.1016/j.cmet.2016.09.001

64. Lee I, Mitsumoto H, Lee S, et al. Higher Glycemic Index and Glycemic Load Diet Is Associated with Slower Disease Progression in Amyotrophic Lateral Sclerosis. Ann Neurol. Feb 2024;95(2):217–229. doi:10.1002/ana.26825

65. Ma Y, Liu Z, Deng L, et al. FGF21 attenuates neuroinflammation following subarachnoid hemorrhage through promoting mitophagy and inhibiting the cGAS-STING pathway. J Transl Med. 2024/05/08 2024;22(1):436. doi:10.1186/s12967-024-05239-y

66. Yang L, Nao J. Focus on Alzheimer’s Disease: The Role of Fibroblast Growth Factor 21 and Autophagy. Neuroscience. Feb 10 2023;511:13–28. doi:10.1016/j.neuroscience.2022.11.003

67. Kakoty V, K CS, Tang RD, Yang CH, Dubey SK, Taliyan R. Fibroblast growth factor 21 and autophagy: A complex interplay in Parkinson disease. Biomed Pharmacother. Jul 2020;127:110145. doi:10.1016/j.biopha.2020.110145

68. Sun Y, Wang Y, Chen ST, et al. Modulation of the Astrocyte-Neuron Lactate Shuttle System contributes to Neuroprotective action of Fibroblast Growth Factor 21. Theranostics. 2020;10(18):8430–8445. doi:10.7150/thno.44370

69. Van Damme P, Robberecht W. Clinical implications of recent breakthroughs in amyotrophic lateral sclerosis. Curr Opin Neurol. Oct 2013;26(5):466–72. doi:10.1097/WCO.0b013e328364c063

70. Paez-Colasante X, Figueroa-Romero C, Sakowski SA, Goutman SA, Feldman EL. Amyotrophic lateral sclerosis: mechanisms and therapeutics in the epigenomic era. Review. Nat Rev Neurol. 05//print 2015;11(5):266–279. doi:10.1038/nrneurol.2015.57

71. Shen Y, Zhu Z, Wang Y, Qian S, Xu C, Zhang B. Fibroblast growth factor-21 alleviates proteasome injury via activation of autophagy flux in Parkinson’s disease. Exp Brain Res. 2024/01/01 2024;242(1):25–32. doi:10.1007/s00221-023-06709-3

72. Teng YD, Mocchetti I, Wrathall JR. Basic and acidic fibroblast growth factors protect spinal motor neurones in vivo after experimental spinal cord injury. Eur J Neurosci. Feb 1998;10(2):798–802. doi:10.1046/j.1460-9568.1998.00100.x

73. Teng YD, Mocchetti I, Taveira-DaSilva AM, Gillis RA, Wrathall JR. Basic fibroblast growth factor increases long-term survival of spinal motor neurons and improves respiratory function after experimental spinal cord injury. J Neurosci. Aug 15 1999;19(16):7037–47. doi:10.1523/jneurosci.19-16-07037.1999

74. Delaye JB, Lanznaster D, Veyrat-Durebex C, et al. Behavioral, Hormonal, Inflammatory, and Metabolic Effects Associated with FGF21-Pathway Activation in an ALS Mouse Model. Neurotherapeutics. Jan 2021;18(1):297–308. doi:10.1007/s13311-020-00933-3

75. Ustyantseva E, Pavlova SV, Malakhova AA, Ustyantsev K, Zakian SM, Medvedev SP. Oxidative stress monitoring in iPSC-derived motor neurons using genetically encoded biosensors of H(2)O(2). Sci Rep. May 27 2022;12(1):8928. doi:10.1038/s41598-022-12807-z

76. Mead RJ, Shan N, Reiser HJ, Marshall F, Shaw PJ. Amyotrophic lateral sclerosis: a neurodegenerative disorder poised for successful therapeutic translation. Nature Reviews Drug Discovery. 2023/03/01 2023;22(3):185–212. doi:10.1038/s41573-022-00612-2

77. Ji K, Zheng J, Lv J, et al. Skeletal muscle increases FGF21 expression in mitochondrial disorders to compensate for energy metabolic insufficiency by activating the mTOR-YY1-PGC1α pathway. Free radical biology & medicine. Jul 2015;84:161–170. doi:10.1016/j.freeradbiomed.2015.03.020

78. Liu X, Wang Y, Hou L, Xiong Y, Zhao S. Fibroblast Growth Factor 21 (FGF21) Promotes Formation of Aerobic Myofibers via the FGF21-SIRT1-AMPK-PGC1α Pathway. Journal of Cellular Physiology. 2017;232(7):1893–1906. 10.1002/jcp.25735

79. Ribas F, Villarroya J, Hondares E, Giralt M, Villarroya F. FGF21 expression and release in muscle cells: involvement of MyoD and regulation by mitochondria-driven signalling. Biochem J. Oct 15 2014;463(2):191–9. doi:10.1042/bj20140403

80. Scaramozza A, Marchese V, Papa V, et al. Skeletal muscle satellite cells in amyotrophic lateral sclerosis. Ultrastructural pathology. Oct 2014;38(5):295–302. doi:10.3109/01913123.2014.937842

81. Manzano R, Toivonen JM, Calvo AC, et al. Altered in vitro Proliferation of Mouse SOD1-G93A Skeletal Muscle Satellite Cells. Neurodegenerative Diseases. 2012;11(3):153–164. doi:10.1159/000338061

82. Martini M, Dobrowolny G, Aucello M, Musarò A. Postmitotic Expression of SOD1^G93A^ Gene Affects the Identity of Myogenic Cells and Inhibits Myoblasts Differentiation. Mediators Inflamm. 2015/09/28 2015;2015:537853. doi:10.1155/2015/537853

83. Allodi I, Montanana-Rosell R, Selvan R, Low P, Kiehn O. Locomotor deficits in a mouse model of ALS are paralleled by loss of V1-interneuron connections onto fast motor neurons. Nat Commun. May 31 2021;12(1):3251. doi:10.1038/s41467-021-23224-7

84. Frey D, Schneider C, Xu L, Borg J, Spooren W, Caroni P. Early and selective loss of neuromuscular synapse subtypes with low sprouting competence in motoneuron diseases. J Neurosci. Apr 1 2000;20(7):2534–42. doi:10.1523/JNEUROSCI.20-07-02534.2000

85. Mishra M, Wu J, Kane AE, Howlett SE. The intersection of frailty and metabolism. Cell metabolism. May 7 2024;36(5):893–911. doi:10.1016/j.cmet.2024.03.012

86. Yan J, Nie Y, Cao J, et al. The Roles and Pharmacological Effects of FGF21 in Preventing Aging-Associated Metabolic Diseases. Front Cardiovasc Med. 2021;8:655575. doi:10.3389/fcvm.2021.655575

87. Cardoso AL, Fernandes A, Aguilar-Pimentel JA, et al. Towards frailty biomarkers: Candidates from genes and pathways regulated in aging and age-related diseases. Ageing research reviews. Nov 2018;47:214–277. doi:10.1016/j.arr.2018.07.004

88. Pandya VA, Patani R. Decoding the relationship between ageing and amyotrophic lateral sclerosis: a cellular perspective. Brain. Apr 1 2020;143(4):1057–1072. doi:10.1093/brain/awz360

89. Cortes-Araya Y, Stenhouse C, Salavati M, et al. KLB dysregulation mediates disrupted muscle development in intrauterine growth restriction. J Physiol. Apr 2022;600(7):1771–1790. doi:10.1113/jp281647

90. Larson KR, Jayakrishnan D, Soto Sauza KA, et al. FGF21 Induces Skeletal Muscle Atrophy and Increases Amino Acids in Female Mice: A Potential Role for Glucocorticoids. Endocrinology. Jan 16 2024;165(3)doi:10.1210/endocr/bqae004

91. Oost LJ, Kustermann M, Armani A, Blaauw B, Romanello V. Fibroblast growth factor 21 controls mitophagy and muscle mass. Journal of cachexia, sarcopenia and muscle. Jun 2019;10(3):630–642. doi:10.1002/jcsm.12409

92. Tezze C, Romanello V, Desbats MA, et al. Age-Associated Loss of OPA1 in Muscle Impacts Muscle Mass, Metabolic Homeostasis, Systemic Inflammation, and Epithelial Senescence. Cell metabolism. Jun 6 2017;25(6):1374–1389.e6. doi:10.1016/j.cmet.2017.04.021

