## Supplementary Materials for "The myokine FGF21 associates with enhanced survival in ALS and mitigates stress-induced cytotoxicity"

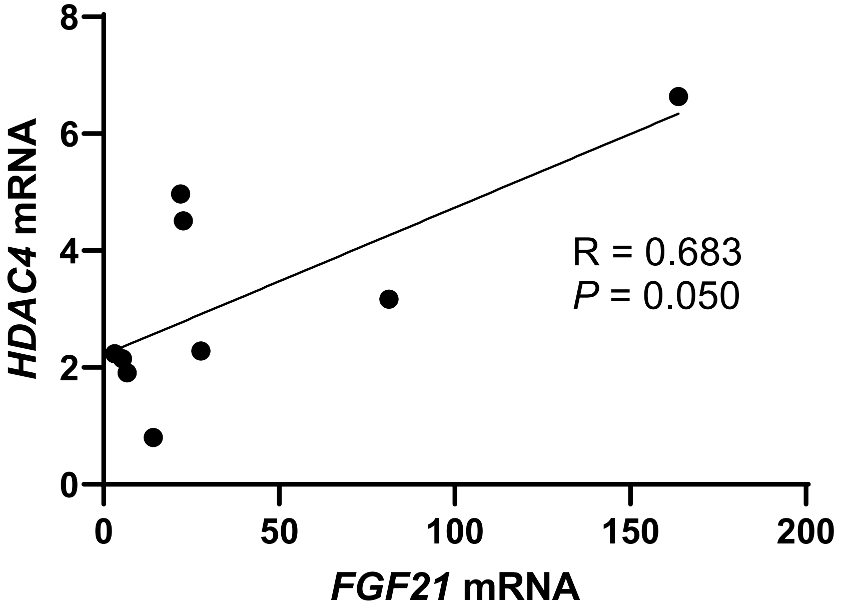


**Supplementary Figure 1.** Spearman rank test correlation between *HDAC4* and *FGF21* mRNA expression levels (assessed by qPCR) in 9 post-mortem ALS muscle samples.

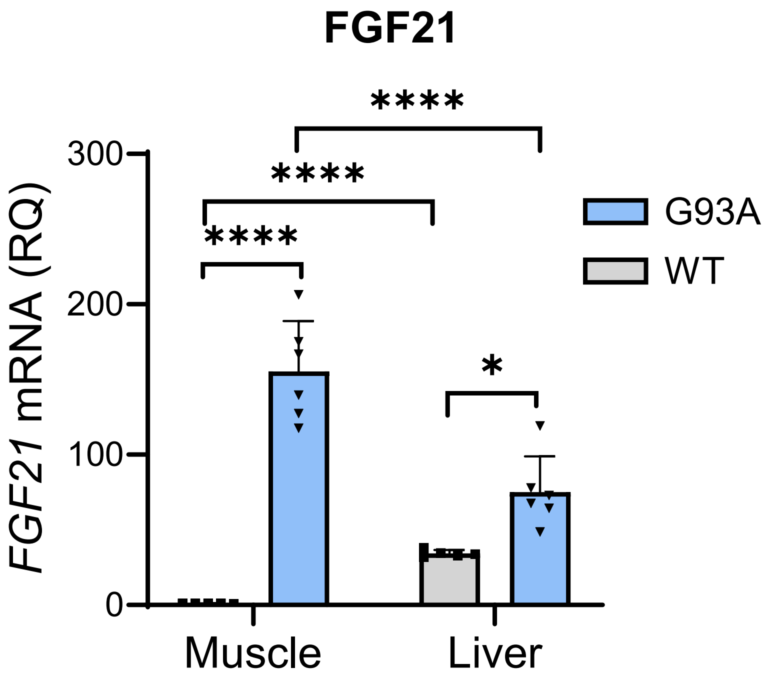


**Supplementary Figure 2. FGF21 mRNA is increased in muscle and liver tissue in the SOD1$^{G93A}$ mouse.** RNA was extracted from wild-type (WT) and SOD1$^{G93A}$ mouse tissue at post-natal day 60 and assessed by qPCR for FGF21 mRNA. All values represent fold-change compared to WT muscle which was set at 1. Data points represent individual mice and bars represent the mean ± SD. *$P < 0.05$, ****$P < 0.0001$; one-way ANOVA followed by Tukey's multiple comparisons test.

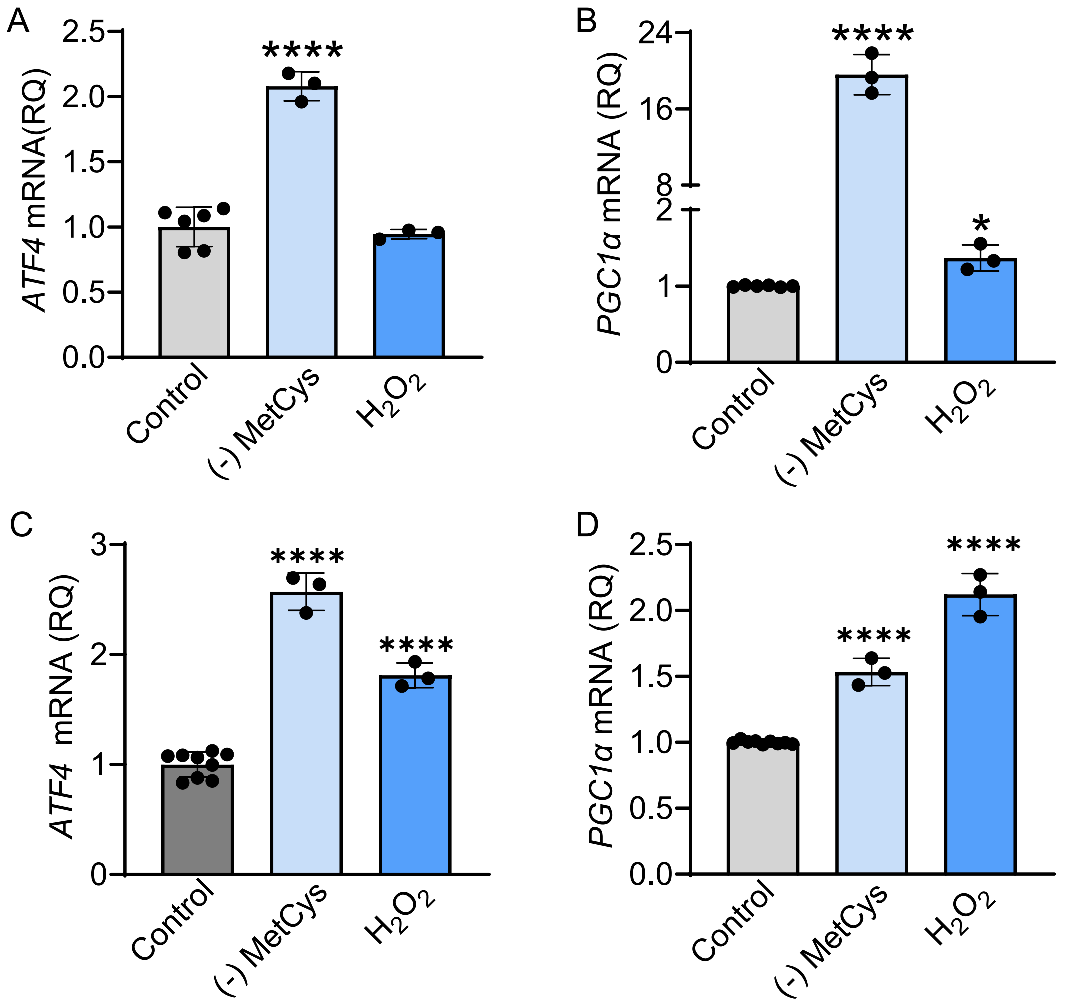


**Supplementary Figure 3. Oxidative stressors induce *ATF4* and *PGC-1α* in NSC-34 motor neuron-like cells and C2C12 myoblasts**. (A) and (B) *ATF4* and *PGC-1α* mRNA levels in NSC-34 cells were quantified after treatment with 100 μM H2O2 or methionine-cysteine (MetCys)-depleted media for 24 h. (C) and (D) *ATF4* and *PGC-1α* mRNA levels were quantified in C2C12 myoblasts after exposure to the same conditions as in (A) and (B). Data points are independent biological samples and bars represent the mean ± SD.*$P$ = 0.048, ****$P < 0.0001$; one-way ANOVA followed by Tukey's multiple comparisons test.

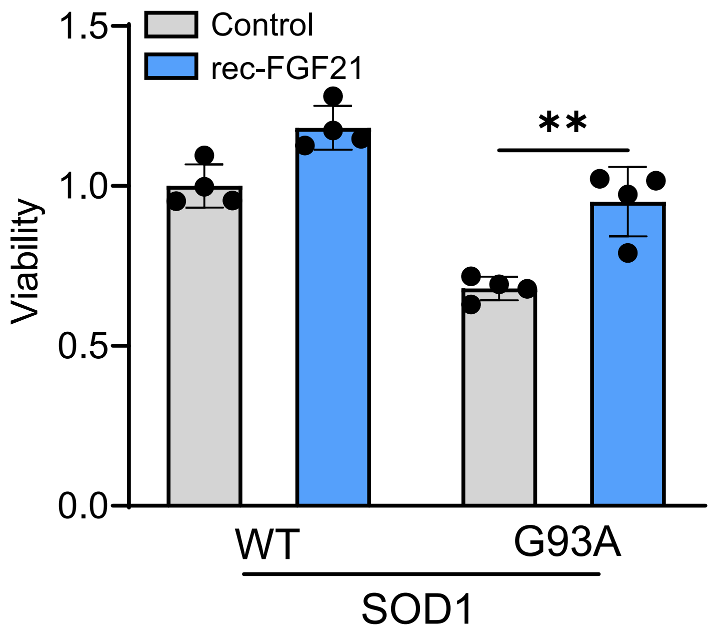


**Supplementary Figure 4. Recombinant FGF21 treatment reverses cytotoxicity of NSC-34 motor neuron-like cells expressing SOD1$^{G93A}$.** NSC-34 cells expressing either WT SOD1 or SOD1$^{G93A}$ were treated with recombinant FGF21 (100 ng/ml). Cell viability was measured 24 hours later as described in the methods. Bars represent the mean ± SD of 4 independent biological samples. **P = 0.003; unpaired two-tailed t-test.

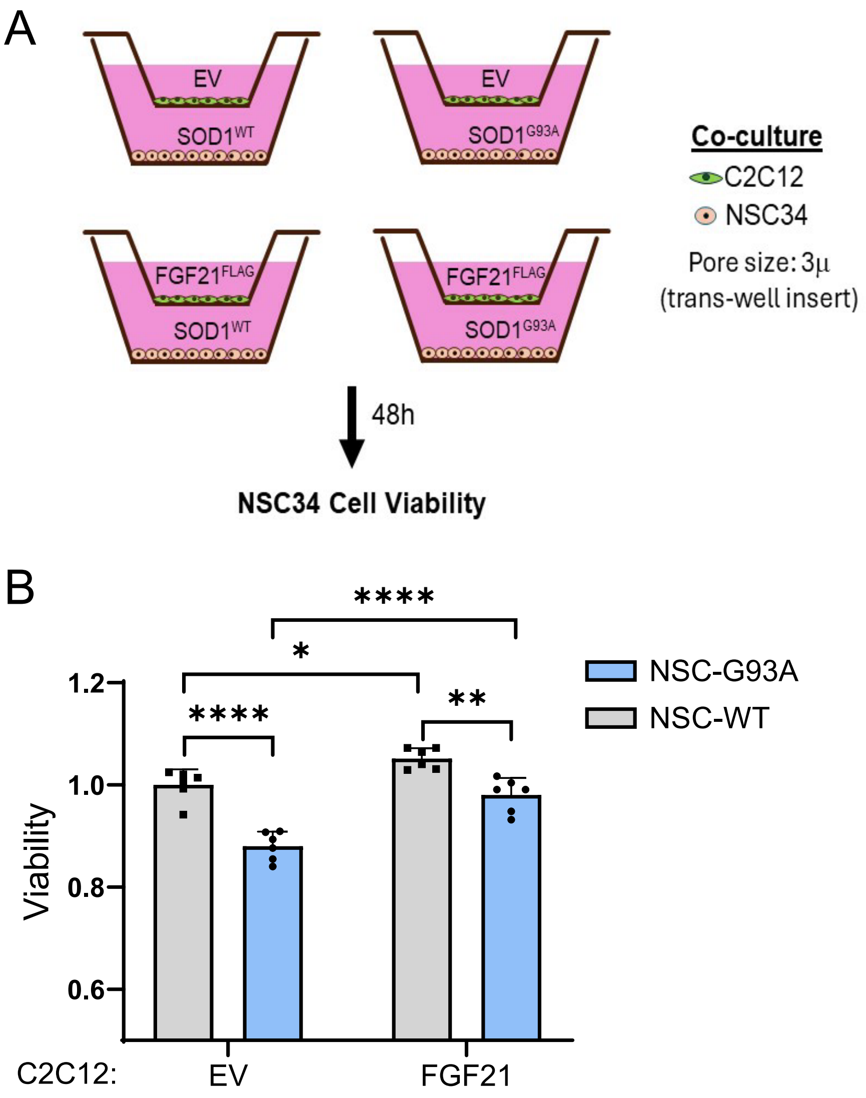


**Supplementary Figure 5. C2C12 cells expressing FGF21 rescue SOD1$^{G93A}$-mediated toxicity in NSC-34 motor neuron like cells in co-culture.** (A) Schematic of co-culture system used. (B) Cell viability of NSC-34 cells expressing either WT-SOD1 or SOD1$^{G93A}$ (lower well) was assessed in the presence or absence of FGF21-expressing C2C12 cells (upper well). Data points represent biological replicates and bars are the mean ± SD. *$P$ = 0.025, **$P$ = 0.002, ****$P$ < 0.0001; one-way ANOVA followed by Tukey's multiple comparisons test.

**Supplementary Table 1: Demographic and clinical data for iPSC-derived motor neurons.**

| Cell Line | Sex | Age (y) | Clinical Diagnosis | Primary tissue | Mutation |
|---|---|---|---|---|---|
| **CS0002iCTR** | M | 51 | Normal | PBMC | N/A |
| **CS83iCTR** | F | 21 | Normal | Fibroblast | N/A |
| **CS188iCTR** | M | 80 | Normal | PBMC | N/A |
| **CS14iCTR** | F | 35 | Normal | Fibroblast | N/A |
| **CS00iCTR** | M | 6 | Normal | Fibroblast | N/A |
| **FA0000011** | F | 49 | Normal | Fibroblast | N/A |
| **NN0003920** | M | 64 | Normal | Fibroblast | N/A |
| **CS0118iALS-SOD1-I114T** | F | 73 | ALS | Fibroblast | SOD1 I113T |
| **CS28iALS** | M | 47 | ALS | Fibroblast | C9ORF72 (HRE ~800) |
| **CS29iALS** | M | 47 | ALS | Fibroblast | C9ORF72 (HRE ~800) |
| **CS52iALS** | M | 49 | ALS | Fibroblast | C9ORF72 (HRE ~800) |
| **CS30iALS** | F | 51 | ALS | Fibroblast | C9ORF72 (HRE ~70) |
| **NN0004306** | F | 51 | ALS | Fibroblast F10330 | C9ORF72 (HRE 2.7kb) |
| **NN0004307** | M | 57 | ALS | Fibroblast F09152 | C9ORF72 (HRE 6-8kb) |

ALS = amyotrophic lateral sclerosis; F = female; HRE = hexanucleotide repeat expansion; M = male; ORF = open reading frame; PBMC = peripheral blood mononuclear cells; SOD1 = superoxide dismutase 1; y = years

**Supplementary Table 2. Demographic and clinical data of tissue samples.**

| | Biopsy | | Autopsy | |
|---|---|---|---|---|
| | **Normal** | **ALS** | **Normal** | **ALS** |
| **Number** | 24 | 36 | 22 | 23 |
| **Mean age (years)** [a] | 52 ± 15 | 57 ± 13 | 67 ± 12 | 64 ± 11 |
| **Age range (years)** | 24 - 77 | 27- 86 | 34 - 83 | 40 - 81 |
| **Gender (M:F)** | 11:13 | 21:15 | 18:4 | 18:5 |
| **Duration** [b] **(m)** | | 15 ± 9 | | 51 ± 32 |
| **Diagnosis** | | Spinal onset (33 )<br>Bulbar onset (3) | | Spinal onset (20 )<br>Bulbar onset (3) |
| **Muscle sampled** | | | | |
| **Biceps brachii** | 5 | 2 | | 2 |
| **Deltoid** | 3 | 11 | 4 | 3 |
| **Vastus lateralis** | 15 | 8 | 3 | 5 |
| **Tibialis anterior** | 1 | 15 | | Triceps (1) |

[a] Mean age (± SD) at time of sample collection.
[b] Mean duration (± SD) from onset of symptoms to sample collection. Duration was unknown for three ALS patients in the biopsy pool.

### Supplementary Table 3: Demographic and clinical data of plasma samples

| | Normal | ALS |
|---|---|---|
| Number | 23 | 28 |
| Mean age (y) [a] | 61 ± 9 | 59 ± 10 |
| Age range (y) | 45 - 84 | 35- 82 |
| Gender (M:F) | 12:11 | 18:10 |
| Duration [b] (m) | | 26 ± 20 |
| Onset | | Spinal onset (22) |
| | | Bulbar onset (6) |

F = female; M = male; m = months; y = years

[a] Mean age (± SD) at time of sample collection.

[b] Mean duration (± SD) from onset of symptoms to sample collection.

### Supplementary Table 4: ALS study patients

| **Plasma FGF21 (FC)[a]** | **< 1.5** | **≥ 1.5** |
|---|---|---|
| Number | 7 | 9 |
| Age (y)[b] | 65 ± 9 | 57 ± 9[d] |
| Age range (y) | 58 - 82 | 42 – 70[d] |
| Gender (M:F) | 4:3 | 7:2 |
| Duration[c] (m) | 18 ± 10 | 26 ± 23[d] |
| Onset | Bulbar (4)<br>Spinal (3) | Spinal (9) |

F = female; FC = fold-change; M = male; m = months; y = years

[a]Fold-change over normal control group

[b]Mean age (± SD) at time of sample collection.

[c] Mean duration (± SD) from onset of symptoms to sample collection.

[d]No significant difference between the 2 groups
